# Predicting base editing outcomes using position-specific sequence determinants

**DOI:** 10.1101/2021.09.16.460622

**Authors:** Ananth Pallaseni, Elin Madli Peets, Jonas Koeppel, Juliane Weller, Luca Crepaldi, Felicity Allen, Leopold Parts

**Affiliations:** Wellcome Sanger Institute, Wellcome Genome Campus, Hinxton, UK; Department of Computer Science, University of Tartu, Tartu, Estonia

**Author notes:** These authors contributed equally to this work (Ananth Pallaseni, Elin Madli Peets).

## Abstract

Nucleotide-level control over DNA sequences is poised to power functional genomics studies and lead to new therapeutics. CRISPR/Cas base editors promise to achieve this ability, but the determinants of their activity remain incompletely understood. We measured base editing frequencies in two human cell lines for two cytosine and two adenine base editors at ∼14,000 target sequences. Base editing activity is sequence-biased, with largest effects from nucleotides flanking the target base, and is correlated with measures of Cas9 guide RNA efficiency. Whether a base is edited depends strongly on the combination of its position in the target and the preceding base, with a preceding thymine in both editor types leading to a wider editing window, while a preceding guanine in cytosine editors and preceding adenine in adenine editors to a narrower one. The impact of features on editing rate depends on the position, with guide RNA efficacy mainly influencing bases around the centre of the window, and sequence biases away from it. We use these observations to train a machine learning model to predict editing activity per position for both adenine and cytosine editors, with accuracy ranging from 0.49 to 0.72 between editors, and with better generalization performance across datasets than existing tools. We demonstrate the usefulness of our model by predicting the efficacy of potential disease mutation correcting guides, and find that most of them suffer from more unwanted editing than corrected outcomes. This work unravels the position-specificity of base editing biases, and provides a solution to account for them, thus allowing more efficient planning of base edits in experimental and therapeutic contexts.

## Introduction

The CRISPR/Cas toolkit has enabled increasingly fine control over DNA sequences [1]. This technology has already uncovered myriad findings in basic research, identified new cancer targets, and offered novel therapeutic avenues for genetic disorders [2–5]. However, the limitation of generating only insertions and deletions without templated repair, and the stochasticity of outcomes have motivated the development of alternative effector proteins such as base editors [6–10] for more precise genome manipulation.

Base editors reduce the range of mutations generated by Cas9 to primarily base substitutions, and alter DNA without potentially apoptosis-inducing double strand breaks. They consist of a catalytically inactive Cas9 fused to a deaminase and domains that modulate the DNA repair pathways [8,9]. There are two main classes of base editor: adenine base editors that convert adenines into guanines using an adenosine deaminase, and cytosine base editors that convert cytosines into thymines using a cytidine deaminase [11]. Once at the target determined by the gRNA, the base editor deaminates suitable nucleotides, which are then converted to another base via DNA repair. The original reports of feasibility [8,9] have been built on to develop increasingly precise and active enzymes [11–16], and also to expand to C to G editing [17,18].

While powerful, base editors have variable efficacy across loci and within the target [8,9,19,20], as well as unintended activity [15,19–24]. The window of activity for the most popular base editors is in positions 4 to 8 of the target sequence [8,9,19,20] (“canonical window”), where the protospacer-adjacent motif is at position 21-23. First reports have attributed some of the variability of base editing efficacy across loci to the APOBEC deaminating domain [25,26], which has a preferred TCW sequence motif, but other sources of variation are less understood. Unintended edits can be frequent, both via off-target editing at unintended locations in the genome [15,21–24], and via bystander editing of bases near the target [19,20,24]. Both types of unintended edits depend on the targeted sequence and position within it [8,9,19,20,22,24]. However, the interplay of position, sequence, and other features that drive variation in editing rate, and could help predict editing outcomes, remains poorly characterized.

Here, we measure the editing frequency of two cytosine base editors (BE4GamRA [27] and FNLS [27]) and two adenine base editors (ABE8e [28] and ABE20m [16]) at thousands of target sites, and uncover new sequence biases that strongly confound the known position-specific editing rates. We use this understanding to build position-specific models of base editing and present a new tool that accurately predicts editing frequency across a range of datasets.

## Results

### Target base context and gRNA efficacy influence editing rate

We set out to quantify the sequence- and gRNA-dependent frequency of editing by cytosine and adenine base editors. We chose two cytosine base editors to screen: FNLS [27], a version of the BE3 editor with an altered nuclear localization signal, and BE4GamRA [27] (hereafter referred to as BE4), an optimization of BE4Gam [29]. We also chose two adenine editors: ABE8e [28] and ABE8.20-m [16] (hereafter referred to as ABE20m), both directed evolutions of ABE7.10 [9] with mutations selected for increased editing efficiency. Following [30], we employed a library of self-targeting constructs which encode both a protospacer adjacent motif-endowed target sequence embedded within 79nt of randomized sequence context, and an expression cassette for a gRNA matching the target. After introducing these constructs into cells, and allowing editing to occur, we sequenced the targets (Figure 1A). We measured base editing frequency in the K562 and HEK293T human cell lines, with a median screen coverage of 890x for cytosine editors and 470x for adenine editors, and sequencing coverage of 1500x (Methods, Supplementary Data). After filtering, we recovered the fraction of edited reads (“editing rate”) for each base of 14,409 target sequences, and observed excellent reproducibility between replicates (Combined Pearson’s R across all positions from 0.73 to 0.91, Figure S1A-D).

**Figure 1.**
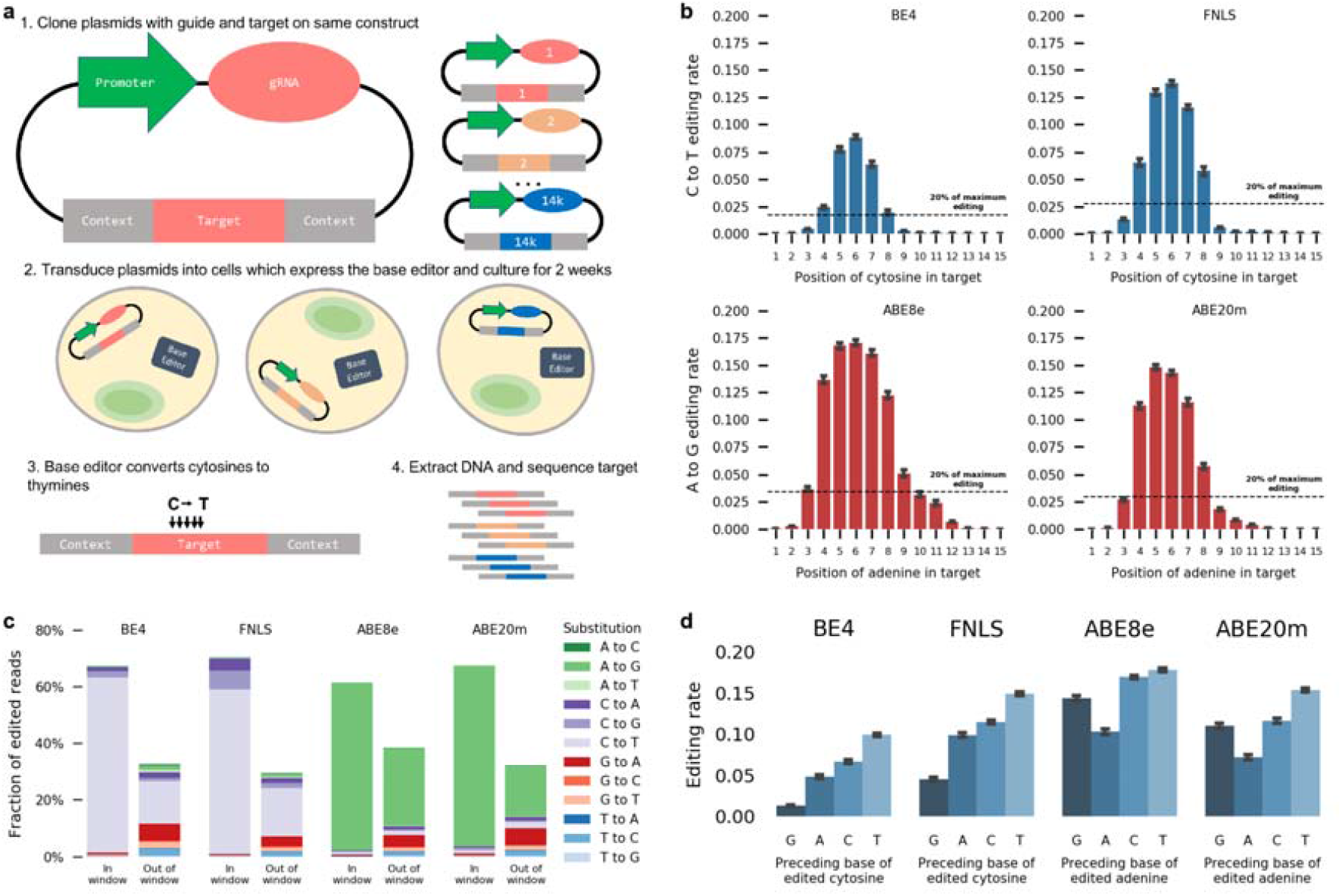
**(a)** A method for high throughput measurement of base editor outcomes. (1) Constructs containing both a gRNA and its target sequence (matched colors) in variable context (gray boxes) are cloned into target vectors containing a human U6 promoter (green). (2) The constructs are packaged into lentiviral particles and used to infect cells that either express base editor protein or have been transfected with base editor protein. (3) The base editors (C to T as an example here) generate base substitutions in the target. (4) DNA from cells is extracted, the target sequence and context are amplified with common primers, and the mutations in the target are determined by sequencing. **(b)** C to T and A to G editing is highest at positions 4-8 in the target sequence. Median C to T (blue bars) or A to G (red bars) editing activity (y-axis) at positions 1-20 in the target sequence (x-axis). Error bars: 95% confidence intervals from 1000 bootstrap samples. Black dashed line: 20% of the maximum editing value found at any position. **(c)** Intended edits in the canonical window are the most frequent outcome for both cytosine and adenine editors. Cumulative frequency (y-axis) of substitution type (color) in the canonical window (position 4-8, left bars) and outside (right bars) for all four editors (x-axis). **(d)** Median base editing rate across targets (y-axis) is influenced by preceding base (x-axis). Error bars: 95% confidence intervals from 1000 bootstrap samples.

We uncovered both known and novel biases in base editing outcomes. The median editing rate across targets was highest at position 6 of the sequence for all editors, and decreased with distance from this position (Figure 1B). Cytosines and adenines in the canonical window (positions 4 to 8) were substantially edited, with rates above 20% of that at position 6, while those outside the canonical window had rates below 10% (Figure 1B). The editing rate per cytosine did not change depending on the number of cytosines in the window for cytosine editors (editing rate per cytosine between 0.04 and 0.05 for BE4, Figure S1E), but decreased with more editable adenines in the window for adenine editors (from 0.13 with only one adenine to 0.08 with six adenines in ABE8e, Figure S1E). When multiple editable bases were present, editing rates were highly correlated for neighbouring bases (average Pearson’s R=0.74 across all editors, Figure S1F), but only moderately correlated otherwise (average Pearson’s R=0.42 across all editors, Figure S1F).

Unintended edits accounted for 35%-42% of all mutated reads across editors (Figure 1C). The most frequent unintended editing event in cytosine editors were C to T edits outside the canonical window (15% and 17% of all mutated reads in BE4 and FNLS, respectively). G to A editing outside the window was the second most common unintended edit in BE4 (6% of all mutated reads) and was relatively frequent in FNLS as well (3.6%), consistent with editing on the opposite strand (Discussion). We also observed frequent transversion edits in cytosine editors, accounting for 6.8% of all mutated reads in BE4 (3.5% C to A, 3.3% C to G) and 14.5% in FNLS (6.2% C to A, 8.3% C to G). The adenine editors had different biases, with the most common unintended edits in both being A to G and G to A outside the canonical window (27% and 4% of all mutated reads in ABE8e, 17% and 5% in Abe20m). Transversion edits were less common in adenine editors, accounting for less than 1% of mutated reads in both ABE8e and ABE20m. All remaining substitutions combined comprised less than 9% of mutated reads in any editor, with less than 2% of the total each (Figure 1C), and their rates outside the window were consistent with those measured in control cells without base editors (Figure S1G). Finally, insertion and deletion frequency in the target window remained below 0.5% (Figure S1H), and we do not consider them further. Overall, nearly two thirds of observed edits were the intended transitions in the canonical window, and the bias towards intended transitions was smaller outside of the window.

Besides position in the targeted sequence, the other known influences of editing frequency are the identity of flanking bases and the gRNA efficacy [8,19,20]. For all editors, the rate of the intended substitution was highest when it was preceded by a thymine (155%, 70%, 25% and 52% increase compared to the other three bases, in BE4, FNLS, ABE8e and ABE20m, respectively; t test p < 10^−20^ in each), consistent with the editing motifs of APOBEC [26] and tadA [31]. C to T editing by cytosine editors was lower when preceded by a guanine (82% and 63% decrease in BE4 and FNLS respectively, t test p < 10^−20^, Figure 1D). A to G editing by adenine editors was lowest when preceded by an adenine (37% and 44% decrease in ABE8e and ABE20m, t test p < 10^−4^, Figure 1D). Editing by all effectors increased when a cytosine followed the edited base and decreased when followed by an adenine. In general, the TNC motif was consistently amongst the best edited sequences.

Sequence identity was important for several unintended substitutions as well (Figure S1I). In particular, C to G transversions by cytosine editors increased over 4-fold at the TCT motif, and were more frequent when the C was followed by a T. This effect was recently used to develop C to G editors elsewhere [32–34]. Furthermore, there is evidence that the cytosine base editors also operate on the opposite strand, as G to A edits were found in much greater quantity in cytosine editors than in controls with only wild-type Cas9 (Figure 1C), and these edits also mirrored motif preferences, albeit with lesser effect on the opposite strand (300% increase in TC to TT in BE4, 50% increase in GA to AA). In summary, the base preceding the edited base has the strongest effect on editing activity for intended transitions, and the base following has an effect on transversions as well.

Measures of Cas9 gRNA efficacy were also informative about base editing efficacy. Predictions from two computational models of gRNA quality (DeepSpCas9 [35] and RuleSet2 [36]), as well as the empirically measured wtCas9 mutation efficiency [30] were correlated with C to T editing frequency in cytosine editors (Pearson’s R=0.13, 0.05 and 0.14, respectively in BE4; 0.11, 0.03, 0.12 for FNLS; p < 0.01 in both) and A to G frequency in adenine editors (Pearson’s R=0.10, 0.11 and 0.09 for ABE8e; 0.11, 0.11 and 0.12 in ABE20m; p < 0.01 in both). The top decile of guides as scored by DeepSpCas9 were edited 70% more frequently than the bottom decile in BE4 (35%, 11% and 20% in FNLS, ABE8e and ABE20m, respectively, Figure S1J). Similarly, the RuleSet2 scores and measured Cas9 mutation efficacies were 27% and 62% higher respectively in the top decile of scores compared to the bottom one in BE4 (13% and 25% in FNLS; 23% and 9% in ABE8; 33% and 24% in ABE20m; Figure S1J). Finally, for better expression from the U6 promoter, the first nucleotide of each guide RNA was changed to a guanine, as has been recommended for genome-wide screens [37]. Targets with a G at position 1 therefore have an improved guide-target match, and this increased editing by 20% over targets that did not start with a G. Thus, we successfully captured independent gRNA- and sequence-dependent biases that affected editing rate, with magnitudes of known effects consistent with existing studies [19,20] (Figure S2E-G).

### Sequence effect depends on edited position

Surprisingly to us, the influence of flanking sequence on editing rate differed substantially across edited positions. For C to T edits using cytosine editors, the preceding guanine and thymine had the strongest marginal effect on editing rate. The detrimental impact of a preceding G was larger away from the canonical window center, with a 49% decrease in median editing at position 6 in BE4, but an 89% decrease at position 9 compared to other preceding bases (35% and 93%, respectively, in FNLS; Figure 2). Similarly, a preceding T resulted in a 45% higher median editing rate at position 6, but a 1900% increase at position 2 (32% and 1800% in FNLS; Figure 2). Preceding thymines had a similar effect on A to G editing for adenine editors, with a 10% increase in editing at position 6 for ABE8e (22% for ABE20m) and 377% increase at position 2 (366% for ABE20m; Figure 2). Cumulatively, 38% of all C to T editing by BE4 across positions 4 to 8 was of the cytosine in the TC dinucleotide, but this increased to 73% for positions not in the 4 to 8 range, where activity was otherwise low (35% and 75% in FNLS; Figure 3B). This strong preference indicates the preceding thymine as a major driver of out-of-window cytosine editing. The preceding T effect was less prominent for the adenine editors, shifting the A to G editing rate from 30% in the window to 47% outside of it in ABE8e (33% and 54% in ABE20m, Figure 3B).

**Figure 2.**
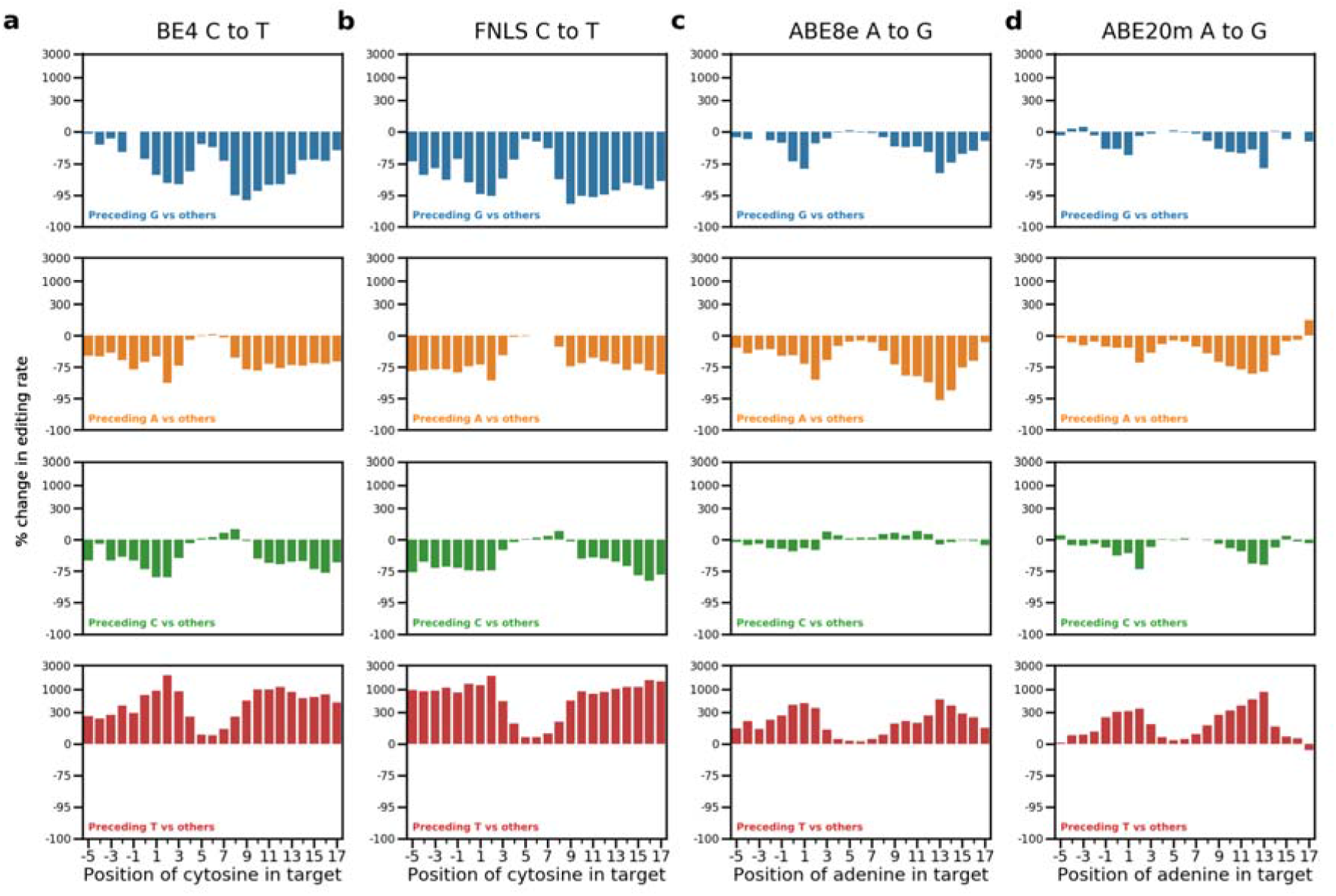
Position-dependent effect of the preceding base. Percentage change in editing rate (y-axis, logarithmic intervals) from having a base preceding the target base (color, rows) compared to all other bases, at different positions in the target sequence (x-axis) for all editors (**a-d**).

**Figure 3.**
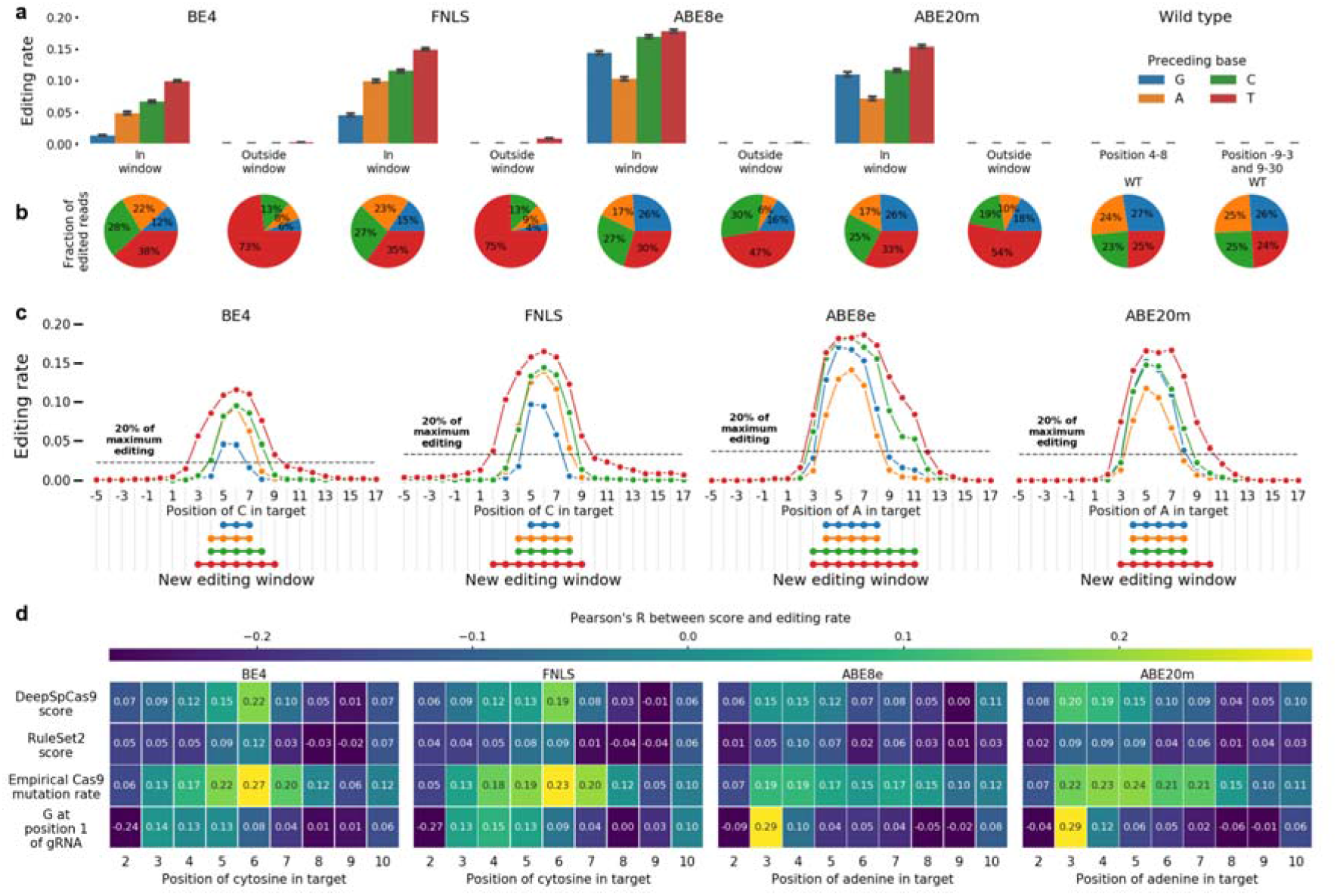
**(a)** Preceding thymines are a driver of out-of-window cytosine editing. Median editing rate across targets (y-axis), for every preceding base (colors), inside and outside canonical target window (x-axis), for cells with base editors (BE4, FNLS, ABE8e, ABE20m) and wild type ones without (WT) compared to the total fraction of C to T or A to G edited reads in the experiment. Canonical window: positions 4-8. Error bars: 95% confidence intervals from 1000 bootstrap samples. **(b)** Same as (a), but fraction of editing outcomes of the total. **(c)** Window of editing changes depending on the base preceding the edited one. Median editing rate (y-axis) of bases at positions -5 to 17 in the target (x-axis) for each preceding base type (colors) for each editor (panels). Black dashed line: 20% of the maximum editing rate at any position for all preceding bases. Linked dots: positions at which editing is above 20% of the maximum. **(d)** Correlation between gRNA quality and editing rate depends on the position. Pearson’s R (color) between measures of gRNA quality (y-axis) and the position of the edited base in the target sequence (x-axis) for each editor (panels).

Position-dependent sequence biases were also present for unintended edits. C to G editing rate in cytosine editors was higher when preceded by a T (88% increase at position 5 and 1186% at position 9 for BE4, S2A, 71% and 1347% for FNLS, Figure S2B), and lower with a preceding G (38% decrease at position 5, 93% decrease at position 9 for BE4, S2A, 28% and 96% for FNLS, Figure S2B). In addition to the preceding base, a following T increased C to G editing across all positions in cytosine editors, with over 115% increase at position 6 in BE4 (Figure S2A). G to A editing is also increased by a following A, consistent with a TC motif on the opposite strand. Altogether, several unintended edits, especially in cytosine editors, exhibited substantial sequence bias that varied across the target (Figures S2A-D).

The large variation in editing rate due to the preceding base suggests a more nuanced redefinition of the cytosine and adenine editing windows. A threshold of 20% of maximum editing for the target produces the canonical window for marginal editing, but gives different editing windows when stratifying by the preceding base. In cytosine editors, the window for cytosines preceded by Cs is positions 4-8, consistent with the canonical window, while a preceding A leads to a window of 4-7. However, with a preceding T, the window broadens to positions 3-9, and a preceding G shrinks it to positions 5-7 (Figure 3C). Similar trends hold for adenine editors, where the A to G window stretched from positions 3 to 11 when adenines were preceded by a T, but was reduced to positions 4 to 8 when preceded by an A or G (Figure 3C). These biases were also present in other large scale measurements of editing rates [19,20] (Figure S2E).

Motivated by the position-dependent sequence effects, we next queried whether the impact of features that capture aspects of gRNA sequence and secondary structure also varies along the target. For cytosine editors, correlation between gRNA features and editing efficacy was strongest at position 6 (Pearson’s R=0.22, 0.12, 0.27 for DeepSpCas9 score, RuleSet2 score and measured Cas9 mutation efficacy respectively in BE4, Figure 3D), but declined with increasing distance from this position. Interestingly, this trend did not hold for the adenine editors, with the largest correlation between A to G editing and the metrics occurring earlier in the sequence for DeepSpCas9 & RuleSet2 scores, or staying consistent across the sequence for the measured Cas9 mutation efficacy (Figure 3D). These effects were also present in other datasets (Figure S2G). Finally, a G at position 1 in the target was associated with increased editing at positions 3 and 4 in all editors and extending to positions 5 and 6 in cytosine editors (Figure 3D). These patterns of feature relevance suggest that gRNA features add bias to the already high editing rates at central positions (especially for cytosine editors), but are less relevant elsewhere. Conversely, sequence bias is lowest centrally, but dominates at outside positions.

### We can accurately predict per-position editing

Given the improved understanding of position-dependent editing rates, we proceeded to build a position-specific editing model. To better generalize across cell types, we combined our data with previously published datasets [19,20], and trained FORECasT-BE, a gradient boosted tree model [38], to predict the normalized editing frequency at positions 3-10 in the target sequence, as well as total fraction of reads edited at any position. Inputs to this model are nucleotide identities at each position in the guide and the melting temperature between the guide and target (Methods). When evaluated on the test set of guides from our experiment only, FORECasT-BE achieved a Pearson’s R of 0.72 across all positions in BE4 (0.71, 0.49, 0.56 in FNLS, ABE8e and ABE20m, respectively, Figure 4A), with highest accuracies at outside positions (Figure S3B). Feature importances in the model reflected the identified sequence biases, with the identity of the base preceding the edited one being most important (Figure S3C-D). We incorporated FORECasT-BE into a command line tool (available at https://github.com/ananth-pallaseni/FORECasT-BE) and a web application (available at https://partslab.sanger.ac.uk/FORECasT-BE), which can be used to predict base editing rates for cytosine and adenine editors.

**Figure 4.**
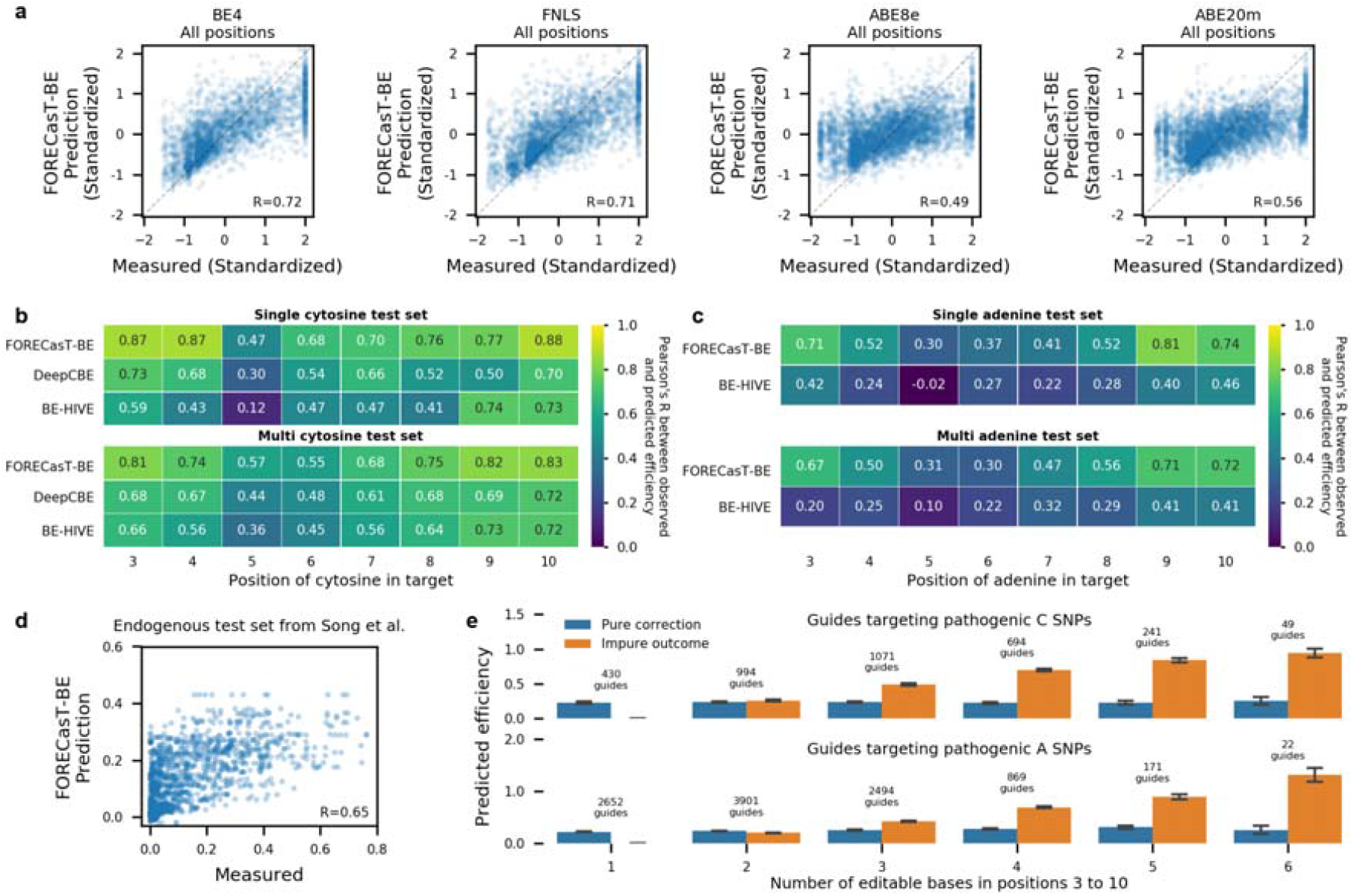
**(a)** FORECasT-BE accurately predicts editing rate. Measured (x-axis) and predicted (y-axis) standardized editing rate (Methods) for guide RNAs (markers) at the different editing window positions (panels) on the multi-cytosine (left panels) or multi-adenine (right panels) test set. Dashed line: y=x. Label: Pearson’s R between measured and predicted scores. **(b-c)** FORECasT-BE outperforms existing prediction models. Pearson’s R between measured and predicted editing rate (color) at different positions in the target (x-axis) for FORECasT-BE and two existing models (y-axis) for cytosine editors (b) and adenine editors (c). Top panel: evaluated on a subset of guides containing only single cytosine in positions 3-10. Bottom panel: evaluated on all guide RNAs. **(d)** FORECasT-BE accurately predicts endogenous editing rate. Measured (x-axis) and predicted (y-axis) standardized editing rate (Methods) for guide RNAs (markers) at the different genomic positions tested in the Song et al. dataset. Dashed line: y=x. Label: Pearson’s R between measured and predicted scores. **(e)** Substantial editing of non-targeted cytosines and adenines for pathogenic corrections. Predicted fraction of impure edits in positions 3-10, (y-axis; orange bars) and correction edits (y-axis; blue bars) for increasing number of editable cytosines (top) or adenines (bottom) in the window (x-axis). Error bars: 95% confidence intervals from 1000 bootstrap samples.

Recently, other methods have been developed for base editing rate prediction. We compared the performance of FORECasT-BE to BE-HIVE [19] that supports both adenine and cytosine editing, and DeepCBE [20], which supports cytosine editing only. These models predict both a total editing rate per guide, as well as a set of specific outcomes, so we evaluate them separately for these tasks. First, we predicted the total fraction of reads mutated across all positions, and observed improved performance of FORECasT-BE relative to other models on our data (Pearson’s R of 0.73 for cytosine editors and 0.77 for adenine editors for FORECasT-BE, 0.60 and 0.60 for BE-HIVE, and 0.60 for DeepCBe on cytosine editors). Next, we restricted evaluation to guides with only a single editable base in the 3-10 window (“single cytosine/adenine test set”), and compared editing rates predicted by FORECasT-BE with outcome predictions (as opposed to total rate predictions) from other models at each position. Our model achieved the highest correlation between measured and predicted cytosine and adenine editing at every position, with lowest improvement over other models at position 6 (Pearson’s R of 0.68, 0.54 and 0.47 for FORECasT-BE, DeepCBE and BE-HIVE respectively for cytosine editors, Figure 4B; 0.41 and 0.22 for FORECasT-BE and BE-HIVE on adenine editors, Figure 4C) and increasing advantage away from it (Pearson’s R of 0.87, 0.73 and 0.59 at position 3 for cytosine editors Figure 4B; 0.71, 0.42 in adenine editors, Figure 4C).

We then performed the same position-specific editing rate comparison on guides with one or more cytosines/adenines in the editing window (“multi-cytosine/adenine test set”). We observed similar patterns across positions (Pearson’s R of 0.55, 0.48 and 0.45 on cytosine editors at position 6, Figure 4B; 0.81, 0.68 and 0.66 for cytosine editors at position 3, Figure 4B; 0.30 and 0.22 on adenine editors for FORECasT-BE and BE-HIVE at position 6, Figure 4C; 0.67 and 0.20 on adenine editors at position 3, Figure 4C). Finally, we considered a combined test set from three studies, and found that while each model performs best on data from the study it was trained on, FORECasT-BE performed well on all of them (at least 90% of the dataset maximum Pearson’s R in all cases, Figure S3E-H), and other models failed to generalize (less than 80% of the dataset maximum Pearson’s R on at least one dataset).

We next tested whether our model generalizes to measurements at endogenous sequences from Song et al. [20], comprising 170-230 guides per editable target position in HCT116 and HEK293T cells. Correlation between predicted and observed editing on this dataset was 0.65 for all editors and positions (Figure 4D), and when stratified by position, ranged from 0.34 in the center to 0.79 at the edges of the window using cytosine editors (0.53 in center and 0.53 at the edges using adenine editors, S3I). This performance was similar to that of other models on the same dataset, with our model improving predictions away from the window center (Figure S3J).

Finally, we used FORECasT-BE to predict point mutation correction purity in disease-relevant contexts. We predicted outcomes at 13,591 pathogenic SNPs from ClinVar which fall within positions 3-10 of a possible target, and can be converted with cytosine or adenine editors [39]. We assumed a 50% conversion rate at position 6, which is the current expectation in a clinical scenario [19], and does not impact the resulting purities. Altogether, 64% of cytosine-targeting guides and 41% of adenine-targeting ones were predicted to have more combined editing of nearby cytosines than the targeted SNP (Figure 4E). The correlation between the expected number of unintended edits and the number of targetable bases in the window was 0.75 for cytosine-targeting guides and 0.76 for adenine-targeting guides, indicating that more targetable bases are expected to produce more unintended edits. While SNP correction purity is influenced by window position and nearby targetable bases (Figure S3K), we identified 108 cytosine-targeting guides and 421 adenine-targeting guides that were predicted to correct the disease-relevant mutations with over 80% purity, and also were in the top 10% of correction efficiencies, and therefore may make reasonable therapeutic targets. For sites where this is not the case, our model can help predict the most frequent unintended edits, which can then be checked for their effect on the protein sequence.

## Discussion

We reported strong position-dependent biases in the determinants of base editing rates from a large survey of editing outcomes in novel and published datasets of cytosine and adenine base editors. Our findings indicate a nuanced action of cytosine editing. The editing window depends on the preceding base, with TC or TA dinucleotides edited beyond the canonical positions 4 to 8. Transversion edits were moderately frequent, and also position-dependent with a strong bias from the following base. In general, as much as 30% of overall editing on average is either out-of-window or not the intended substitution, with out-of-window edits more frequent if following thymines. All the findings on position dependence also held in existing large-scale datasets that were generated using different base editor proteins, cell lines, and target libraries.

We incorporated these insights into a predictive model, FORECasT-BE, trained on multiple datasets to robustly predict cytosine and adenine editing, and found it to be the most accurate model for edits in our K562 and HEK293T data. In comparing models trained on different datasets, we found that diverse training data, as used for FORECasT-BE, improves generalization performance on other data as well. The dataset specificities could be driven by a wide complement of factors ranging from cell type and repair activity to the effector protein used, its delivery mechanism, the guide RNA expression method, etc., which require further study with richer data, and including endogenous and *in vivo* contexts. Models that generalize beyond a single dataset, such as FORECasT-BE will prove a useful tool when more bespoke ones are not available.

Both existing and our new models achieved more accurate predictions at positions further away from the canonical window center. Editing at central positions in our data was as reproducible as for outside positions (Figure S1A-D), so increased measurement noise is unlikely to be the cause. Taken in conjunction with the increased correlation between gRNA-level features and editing at central positions, it is likely that these patterns arise from different strengths of sequence effect on mutation generation, where the target base context has a large influence outside the window and the gRNA features matter more at the center.

Using FORECasT-BE, we find that most pathogenic SNPs that could be reverted using cytosine base editors are expected to have more unintended edits than clean conversions, but identify 108 guides for C to T editing and 421 guides for A to G editing with promise to cleanly correct their targets. The editing of non-targeted cytosines presents a hurdle for clinical use of base editors. While the technology develops to address these issues, predictive models will remain essential to identify unintentional edits in advance, to potentially account for their effects [40], and to evaluate the pathogenicity of a guide when planning therapies.

As base editors are already used in genetic screens [41], have demonstrated feasibility in preclinical settings [42–44], and will likely soon advance into clinical trials, there is a need for better understanding of editing determinants and more accurate models of their effects. This work, and predictive outcome models in general, will be necessary to support the use of base editing in scientific and therapeutic applications.

## Author contributions

Designed project: EMP, LC, FA, AP, JK, LP. Performed experiments: EMP, JW, LC. Performed analysis: AP, JK, JW, EMP, FA. Supervised study: LP. Wrote paper: AP, LP, with input from all authors.

## Acknowledgements

A.P, E.M.P, J.K, J.W and L.C were supported by Wellcome (206194). F.A. was supported by a Royal Commission for the Exhibition of 1851 Research Fellowship. L.P. was supported by Wellcome (206194) and the Estonian Centre of Excellence in IT (EXCITE) (TK148).

## Data and code availability

**Sequencing data:** PRJEB12405, see Supplementary Table 2 for sample accessions

**Processed data:** https://doi.org/10.6084/m9.figshare.14460039

**Guides used in this study:** https://doi.org/10.6084/m9.figshare.14460138

**FORECasT-BE code:** https://github.com/ananth-pallaseni/FORECasT-BE

## Methods

### Cell culture and cell line generation

K562 cells were cultured in RPMI and HEK293FT cells were cultured in Advanced DMEM (Gibco). In both cases supplemented with 10% FCS, 2 mM glutamine, 100 U/ml penicillin and 100 mg/ml streptomycin. Cells were cultured at 37°C, 5% CO2. K562 cells endogenously expressing BE4 and FNLS were generated by infecting K562 cells with a lentiviral vector carrying a base editor and puromycin resistance genes (pLenti-BE4GamRA-P2A-Puro, Addgene 112673; pLenti-FNLS-P2A-Puro, Addgene 110841) [27]. Lentivirus was produced and wildtype K562 cells were infected as described below. 24 h later selection with 2 μg/ml puromycin was started. After one week, the selection was stopped and cells were expanded. After 10 days of expansion, cells were treated for an additional 3 days with 0.5 μg/ml puromycin to enrich for cells with desired constructs.

### Lentiviral library

The lentiviral library used in this study was the same one used in Allen et al., 2019 [30]. Briefly, the library uses the pKLV2-U6gRNA5-PGKpuro2ABFP-W [45] backbone and encodes 41,630 gRNA-target constructs. The gRNA is 23 nt (including protospacer-adjacent motif) and the target sequence is 79 nt flanked with PCR priming sites.

### Lentivirus production and titration

Lentivirus was produced using HEK293FT cells that were transfected with Lipofectamine LTX (Invitrogen). 5.4 μg of a lentiviral vector, 5.4 μg of psPax2 (Addgene 12260), 1.2 μg of pMD2.G (Addgene 12259) were mixed in 3 ml Opti-MEM together with 12 μl PLUS reagent and incubated for 5 min at room temperature. 36 μl of the LTX reagent was added and the mix was incubated for another 30 min at room temperature. 3 ml of the transfection mix was then added to 80% confluent cells in 10 ml DMEM media in a 10-cm dish. After 48 h the supernatant was collected and stored at 4°C. Fresh media was added to the cells and harvested 24 h later. The supernatants from both harvests were mixed and centrifuged overnight at 6,000g at 4°C and then for another 2 h at 20,000g. The supernatant was removed and the viral pellets were resuspended in DPBS resulting in 50x concentration of the virus. The virus was stored at -80 °C. The procedure was scaled up accordingly for a larger production of virus.

For virus titration, K562 cells were seeded into a 96-well plate at 5×10^4^ cells/well. Increasing amounts of virus and 8 μg/ml polybrene (hexadimethrine bromide, Sigma) were added to each well. The plate was centrifuged at 1000g for 30 minutes at room temperature. The cells were resuspended and cultured for 72 h before harvesting for FACS analysis. The viral titer was estimated based on BFP+ cells and scaled up for the following screens. Data was analyzed by FlowJo.

### Screening

All cell lines were infected aiming at a multiplicity of infection (MOI) of 0.8 and a coverage of 800x. Each cell line was infected twice and treated as two biological replicates.

For cytosine base editor screens, K562-BE4 and K652-FNLS cells were seeded at a concentration of 1.5×10^5^ cells/ml. Cells were cultured for 27 days and samples were harvested at 3, 6, 10, 14, 17, 21, 24 and 27 days post-infection. Cells were passaged to maintain higher coverage than at the time of infection. At 4 days post-infection, a subsample of cells was harvested for FACS analysis to estimate the MOI based on BFP+ cells. The data was analysed with FlowJo.

For adenine base editor screens, 293FT cells were cultured in media containing 2 μg/ml puromycin for one week to select for infected cells. 6×10^7^ cells were then seeded into six tissue culture dishes with 150 mm diameter in 20 ml media. 24h later the media was refreshed with 15 ml of media. Transfection mixes were prepared in two steps (protocol adapted from [46]). First, 16 ml of Opti-MEM was mixed with 72 μg of base editor encoding plasmid (ABE8e, Addgene #138489 or ABE8.20-m, Addgene #136300), 8 μg of pCS-GFP plasmid and 800μl Plus reagent. Secondly, 16ml Opti-MEM was mixed with 400 μl Lipofectamine 3000 (Invitrogen) and 1600 μl Lipofectamine LTX. The two solutions were mixed together, incubated for 30 minutes at room temperature and 3.2 ml of the transfection mix was transferred to each tissue culture plate. 48h later 15 ml of media was added to cells. After 14h cells were harvested and a subsample of cells were used to check for transfection efficiency via flow cytometry. The data was analysed with FlowJo.

### Sequencing library preparation

Genomic DNA extraction and sequencing library preparation for the main screens were done as described in Allen et al., 2019 [30]. Briefly, to amplify the target sequence from the gDNA, primers P1 and P2 (Table S1) were used with the Q5 Hot Start High-Fidelity 2X Master Mix (NEB). To ensure coverage for each sample, 416 μg of gDNA was used as template and each PCR reaction was run in 50 μl aliquots containing no more than 5 μg DNA. The PCR reaction was column-purified with the QIAquick PCR Purification Kit (Qiagen). Sequencing adaptors and barcodes were added with a second round of PCR using the KAPA HiFi HotStart ReadyMix (Roche), primers P3 and P4 (Table S1) and 1 ng of template DNA. Amplicons were purified with Agencourt AMPure XP beads in 1.2:1 ratio (beads to PCR reaction volume), quantified with the Quant-iT™ High-Sensitivity dsDNA Assay Kit (Invitrogen). The amplicons for the BE4 and FNLS screens were sequenced using a NovaSeq S4 XP and the ABE8e and ABE20m screens used a HiSeq4000.

To measure mismatch rate between guides and targets, we amplified the target region together with the guide sequence from the genomic DNA in one of the cell pools above before editing occurred using primers P5 and P6 (Table S1). The sequencing library was prepared as described before, using 104 μg of gDNA as template for the PCR. Sequencing adaptors and barcodes were added with a second round of PCR using the KAPA HiFi HotStart ReadyMix (Roche), primers P3 and P4 and 1 ng of template DNA. The libraries were sequenced with a HiSeq 2500 using paired end sequencing, such that the forward reads covered the guide region and the reverse reads covered the target.

### Data processing

We assigned reads to guides, and generated outcome profiles using the custom processing pipeline from Allen et al., 2019 [30]. Outcome profiles for a guide are represented by pairs of mutations and the number of guide reads which had that unique mutation. For convenience, we also stored a matrix of the fraction of guide reads containing every possible base substitution at every position in the target sequence (12 substitutions x 79 positions).

To ensure adequate coverage, we first removed guides with less than 100 reads in any sample (timepoint or replicate) from the analysis.

For BE4 and FNLS, we calculated the correlation of C to T editing at positions 4 to 8 between two replicate screens at each timepoint and found that replicates agreed with each other less at the later timepoints (Figure S4B). We speculate that this is due to the toxicity of the editors, an argument that is supported by the decrease in average C to T editing at timepoints 21-27 (Figure S4A). Thus we chose to combine timepoints 10-17 (which were highly correlated, Figure S4B) in our BE4 and FNLS data, by pooling together all the reads assigned to the same guide in each timepoint and treating this as a single screen. We then calculated the between-replicate correlation of C to T editing on the dataset of combined timepoints at positions 4 to 8, and found that replicates were very similar (Median Pearson’s R across positions of 0.87 and 0.91 in BE4 and FNLS, respectively, Figure S1A-B).

For all editors, we chose to combine the replicates using the same method as with timepoints, by adding read counts.

After taking guides common to the relevant timepoints, removing guides with under 100 reads and only retaining guides for which we had guide-target mismatch information, we are left with 14,000 guides.

### Correcting for mismatched guide-target pairs

To correct for recombination during infection of the guide library, which results in guide-target mismatch in some cells, we calculated the guide-target match rate for each guide using data from a long-range PCR on an early time point in one cell pool. The reads were assigned to guides by first checking if the forward read was a direct match to any of our guide sequences and, if matched, that read was assigned to that target. If the read was not a perfect match, we checked if the middle 10 bases of the read matched the middle 10 bases of any of our guides, as this stretch of bases could uniquely identify 82% of our guides. Once the forward PCR reads were assigned to guides, we aligned the expected target of the assigned guide with the sequence of the reverse read using the pairwise2 method from Biopython [47]. Gaps were given a penalty of -1 and extending a gap was given a penalty of -0.1 (pairwise2.align.globalxs(guide1, guide2, -1, -0.1)). Random alignments of different targets in the dataset to the wrong guide all had scores below 41 (Figure S4C), so we set a threshold of 50 to call a match, and any reads with an alignment score under this were considered mismatched from recombination. We calculated the guide-target match rate for a guide as the fraction of matched reads for that guide.

To correct for recombination mismatch, we scaled the total number of reads in the base editing experiment for each guide by its guide-target match rate to get the number of reads that came from matching constructs. The number of reads with edits was left unchanged under the assumption that the constructs matched to be able to create an edit..

### Measures of gRNA efficacy

DeepSpCas9 scores and RuleSet2 scores were calculated using the 20 nucleotides of the target sequence. DeepCas9 scores were obtained from the batch prediction tool offered at http://deepcrispr.info/DeepSpCas9 with default settings, and RuleSet2 scores were computed using the software from [36], also with default settings. Empirically measured Cas9 mutation efficacy was obtained from [30], which used the same guide library as this study in K562 cells.

### Other base editing efficiency datasets

Data were downloaded from [20] and [19] and used to calculate editing rates for each substitution at each position for every guide. We used the mES-BE4 and mES-ABE datasets from [19] as the closest match to our editors, and for its high concordance of replicates.

We selected guides which had cytosines or adenines at the specified position and standardized the editing rates by subtracting the mean at that position and dividing by the standard deviation. We then appended our BE4 dataset, our FNLS dataset, the mES-BE4 dataset from [19] and the cytosine dataset from [20] together to get a combined cytosine dataset; then we combined our ABE8e dataset, our ABE20m dataset, the mES-ABE dataset from [19] and the adenine dataset from [20] to get a combined adenine dataset.

### Creating training datasets

We maintained the train/test set distinction provided in the Song et al. data, and randomly partitioned the Arbab et al. data into training and test sets with 90% for training and 10% for testing. We randomly separated the guides in our experiment into a training set consisting of 90% of the guides and a test set consisting of the remaining 10% of guides, such that no guide was present in both sets. In order to allow generalizability across experiments, we standardized real editing rates at each position by subtracting the mean edit rate at that position and dividing by the standard deviation at that position. Guides were converted into feature vectors by first one-hot encoding the 20 nt guide sequence and then appending the melting temperature of the 20 nt guide sequence as calculated using the Biopython [47] MeltingTemp function.

### Modelling editing rate

We trained and evaluated models with one-hot encoded sequence features and different sets of guide efficacy features on the training set, using 5-fold cross validation, to select the final set of features used. We found that all combinations of melting temperature, DeepSpCas9 score and RuleSet2 score produced similar contributions to predictive accuracy and functioned as proxies for each other, so we chose to use melting temperature alone.

We predicted standardized editing rate for base editing efficacy at each position using gradient-boosted trees, as implemented in the Scikit-learn package [48]. One gradient-boosted tree model was trained per position. For each one, we chose to use 100 shallow trees (n_estimators), with maximum depth 4 (max_depth), a minimum of two samples per leaf node (min_samples_leaf) and a learning rate of 0.1. These values were obtained through 5-fold cross-validation on the training set, independently testing values more extreme than the selection in both directions (n_estimators 10 to 1000, max_depth 1 to 10, min_samples_leaf 1 to 50, learning rate 0.001 to 1). After training the final models on the full training set, we evaluated their performance on the test set by calculating the correlation between the predicted standardized rate and the measured standardized rate.

### Comparisons with other models of editing rate

We compare FORECasT-BE to two other models, BE-HIVE [19] and DeepCBE [20]. Predictions for DeepCBE were obtained using the batch prediction tool offered at http://deepcrispr.info/DeepBaseEditor/, while those for BE-HIVE were obtained by running the BE-HIVE scripts from https://github.com/maxwshen/be_predict_efficiency. For BE-HIVE, we specified a mean of 0.5 and a standard deviation of 0.25 as the scaling parameters; these linearly scale the outputs and do not affect the correlations to true editing rates. When using the BE-HIVE model to evaluate guides from the Song et al. study, we padded the given 30nt sequences with As to create the required 50nt input.

### Endogenous data comparison

Editing information on endogenous base editing outcomes was obtained from [20] and editing rates for each substitution at each position for every guide were calculated as above. We predicted the per-position editing rate on guides in this dataset to get standardized predictions, and scaled them back into efficiencies using the per-position means and standard deviations obtained from the Song screening dataset [20].

### Prediction in disease contexts

A set of guides targeting pathogenic SNPs correctable by a C to T or A to G substitution was obtained from [39]. We predicted standardized efficiencies for positions 3-10 in this guide set and scaled them into real efficiencies by assuming a mean of 50% editing at position 6 to match the maximum rate in [19]. We computed the predicted correction efficiency as the rate of C to T or A to G editing at the position of the SNP in the guide, and the expected number of unintended edits as the sum of predicted C to T or A to G editing at other positions.

