## Supplementary Information for "Predicting base editing outcomes using position-specific sequence determinants"

##### Figure S1A

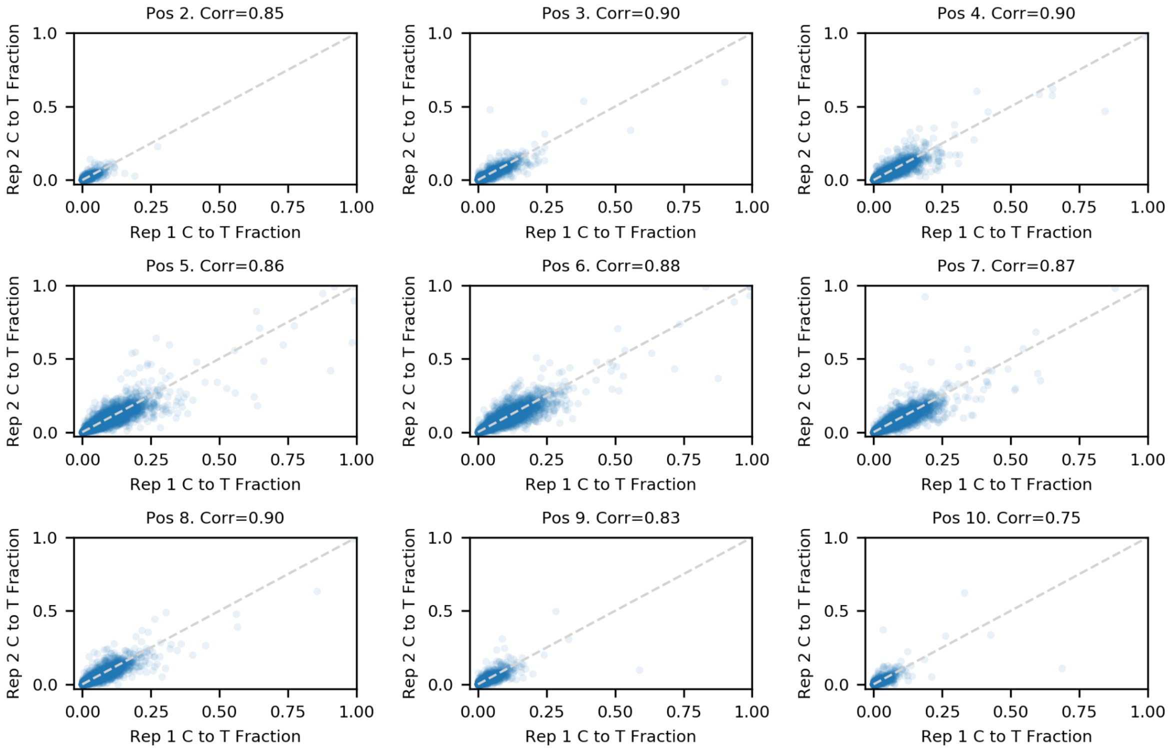


Base editor efficacy is reproducible between replicates in BE4. Fraction of reads with C to T edits in replicate 1 (x-axis) is strongly correlated with the fraction of reads with C to T edits in replicate 2 (y-axis) for cytosine in different positions of the target sequence (panels) across different targets (markers). Dashed line: x=y.

Figure S1B

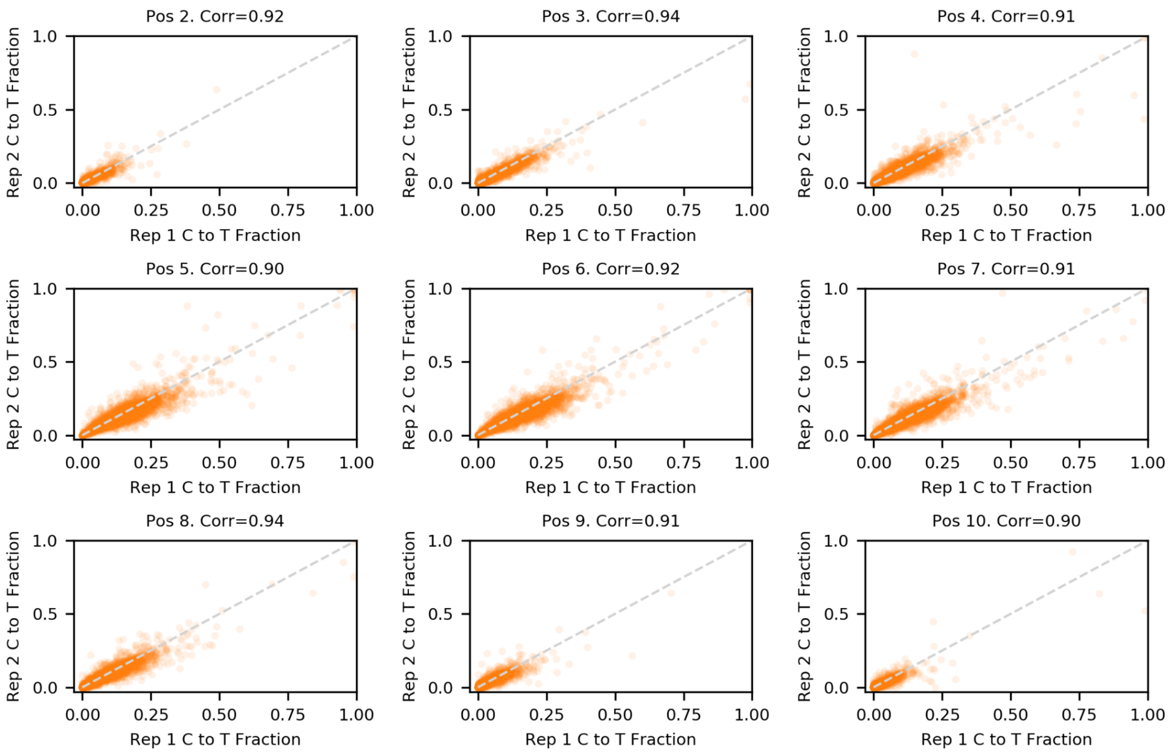


Base editor efficacy is reproducible between replicates in FNLS. Fraction of reads with C to T edits in replicate 1 (x-axis) is strongly correlated with the fraction of reads with C to T edits in replicate 2 (y-axis) for cytosine in different positions of the target sequence (panels) across different targets (markers). Dashed line: x=y.

### Figure S1C

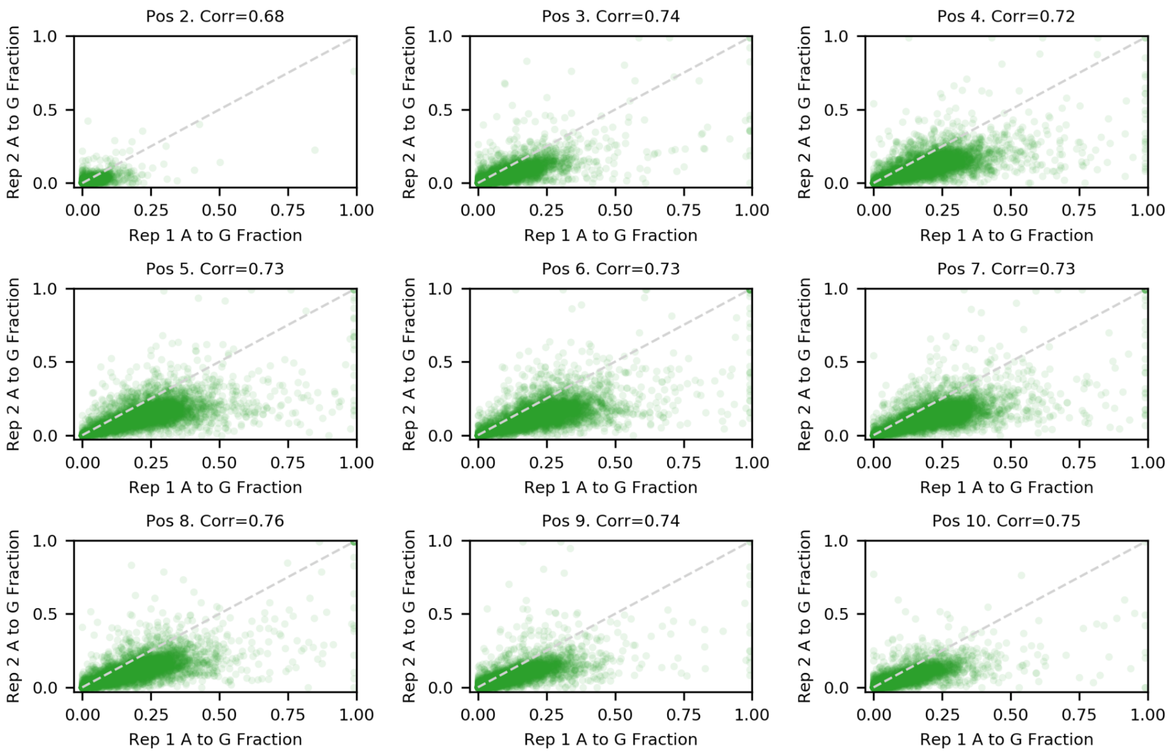


Base editor efficacy is reproducible between replicates in ABE8e. Fraction of reads with A to G edits in replicate 1 (x-axis) is strongly correlated with the fraction of reads with A to G edits in replicate 2 (y-axis) for adenine in different positions of the target sequence (panels) across different targets (markers). Dashed line: x=y.

Figure S1D

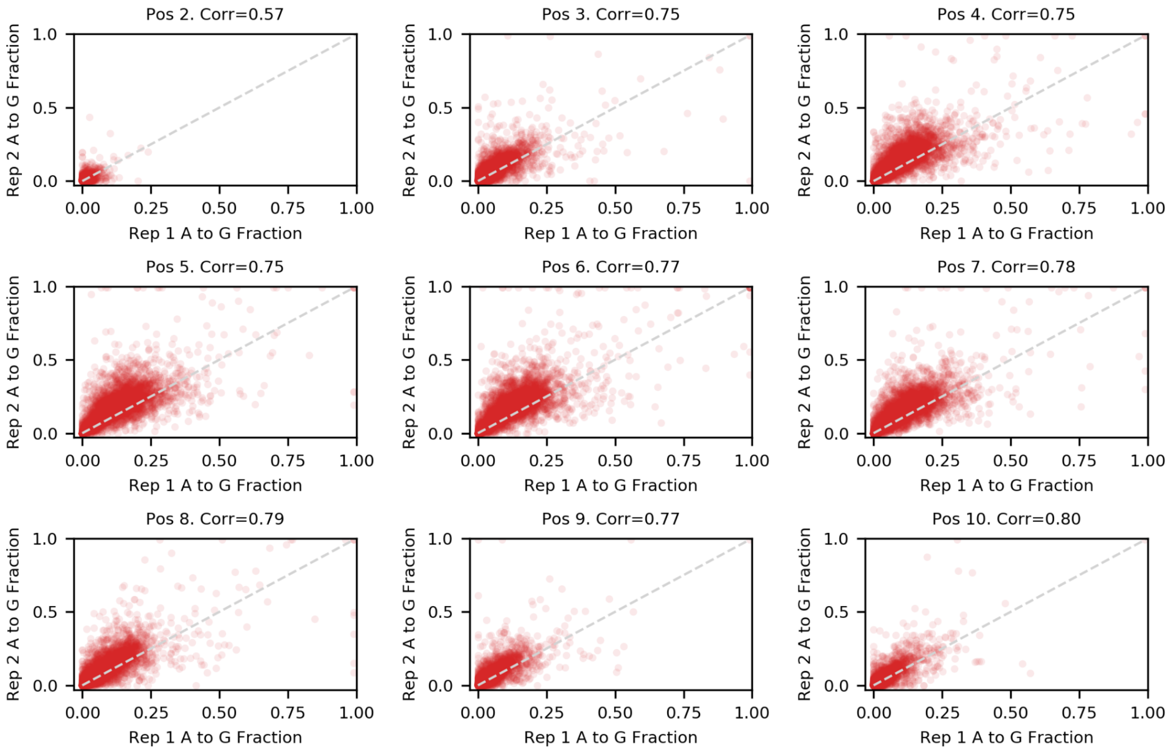


Base editor efficacy is reproducible between replicates in ABE20m. Fraction of reads with A to G edits in replicate 1 (x-axis) is strongly correlated with the fraction of reads with A to G edits in replicate 2 (y-axis) for adenine in different positions of the target sequence (panels) across different targets (markers). Dashed line: x=y.

### Figure S1E

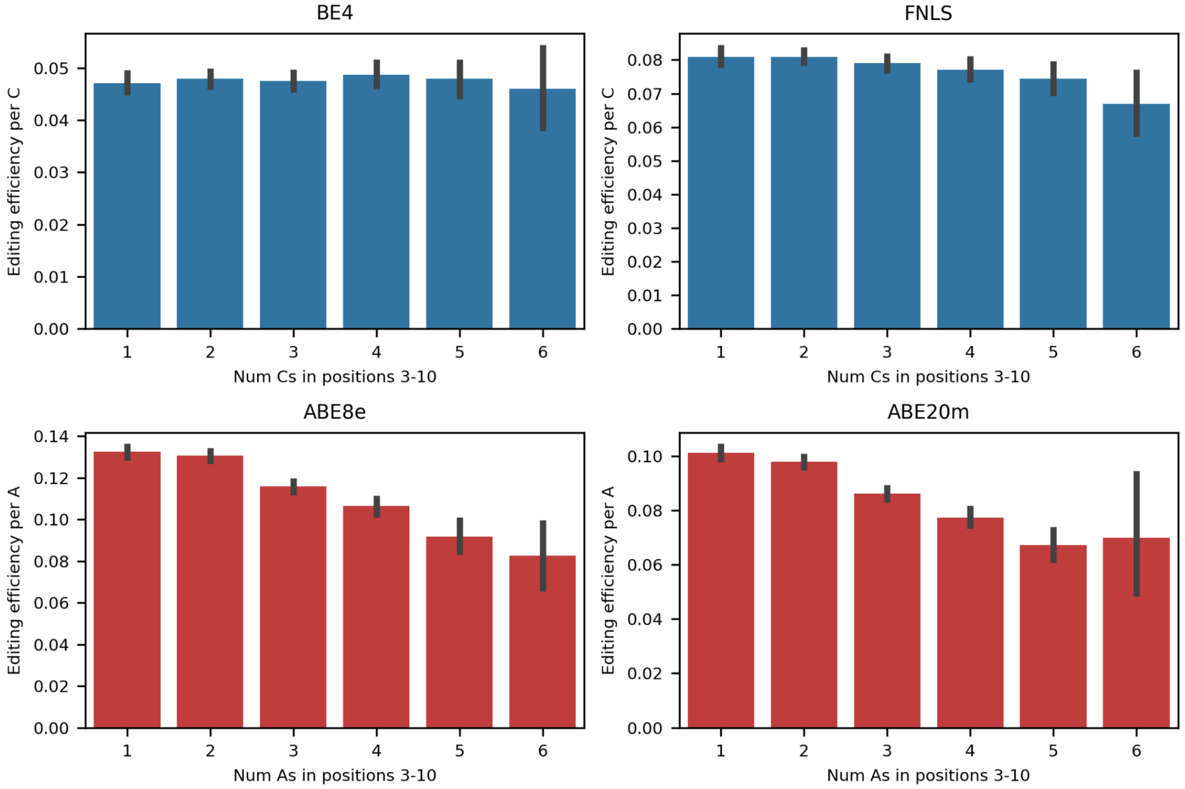


Editing efficiency per number of targetable bases (y-axis) for increasing number of targetable bases in the editing window (x-axis) for BE4 (top left), FNLS (top right), ABE8e (bottom left) and ABE20m (bottom right). Error bars: 95% confidence intervals from 1000 bootstrap samples.

### Figure S1F

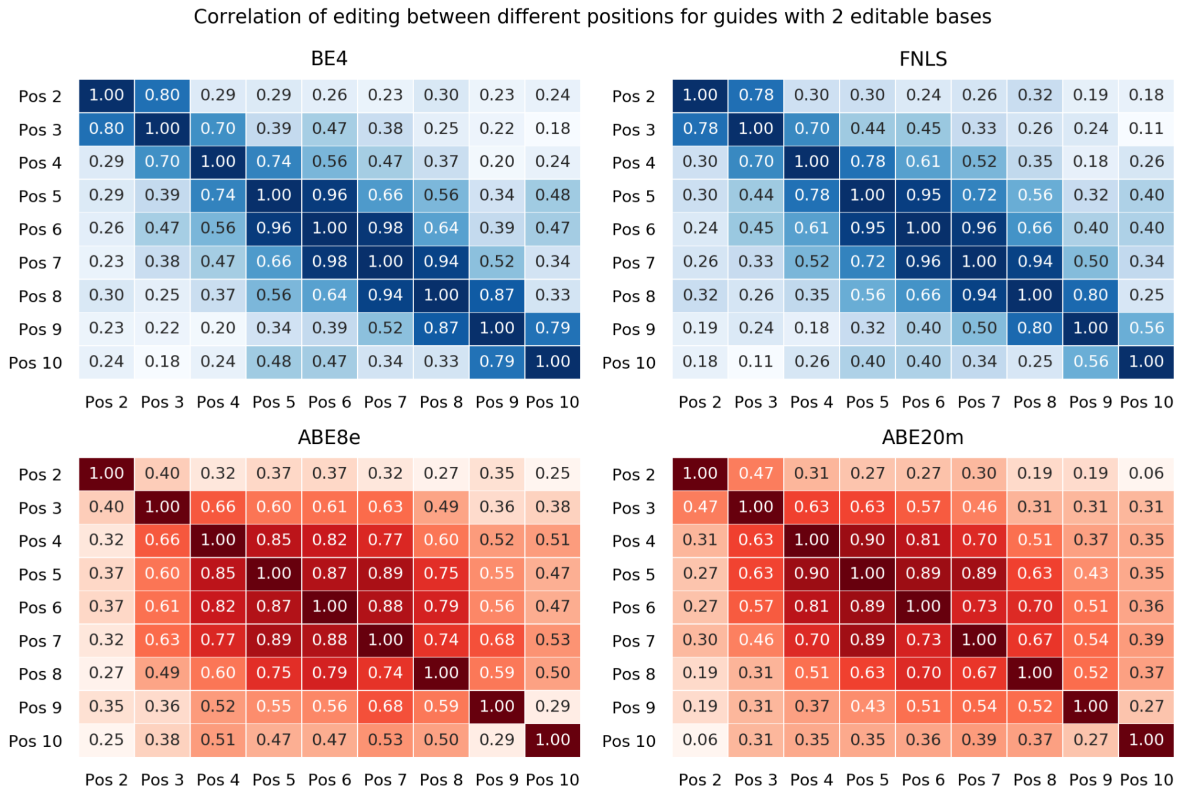


Editing at nearby positions is correlated. Pearson's R between editing at one position (x-axis) and another position (y-axis) for guides with 2 targetable bases in positions 2-10 for BE4 (top left), FNLS (top right), ABE8e (bottom left) and ABE20m (bottom right).

### Figure S1G

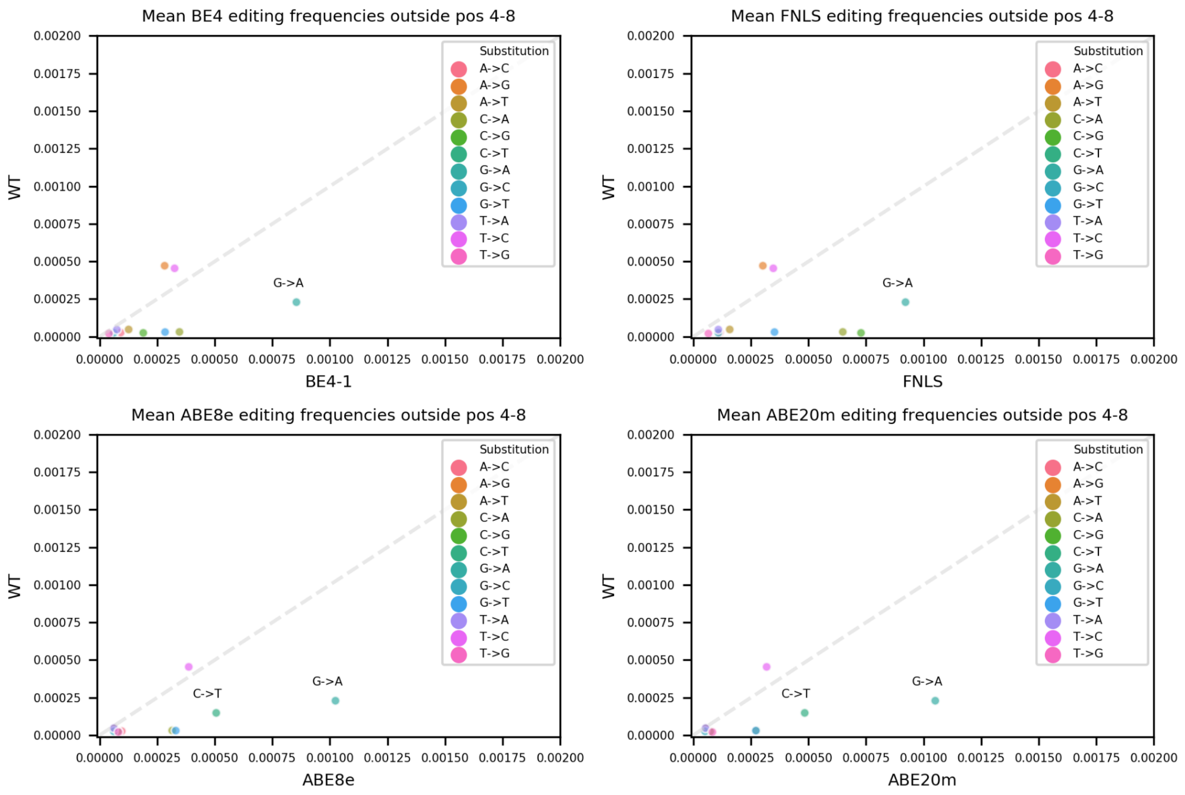


C to T and A to G editing outside positions 4 to 8 predominantly occurs in cells with base editors. Editing rate outside positions 4 to 8 in the target for wild type cells (y-axis) and base editor endowed cells (x-axis) for BE4 (top left), FNLS (top right), ABE8e (bottom left) and ABE20m (bottom right) for different substitution types (colors). Dashed line: y=x. Intended edits (C to T in the top panels, and A to G in the bottom panels) are omitted.

### Figure S1H

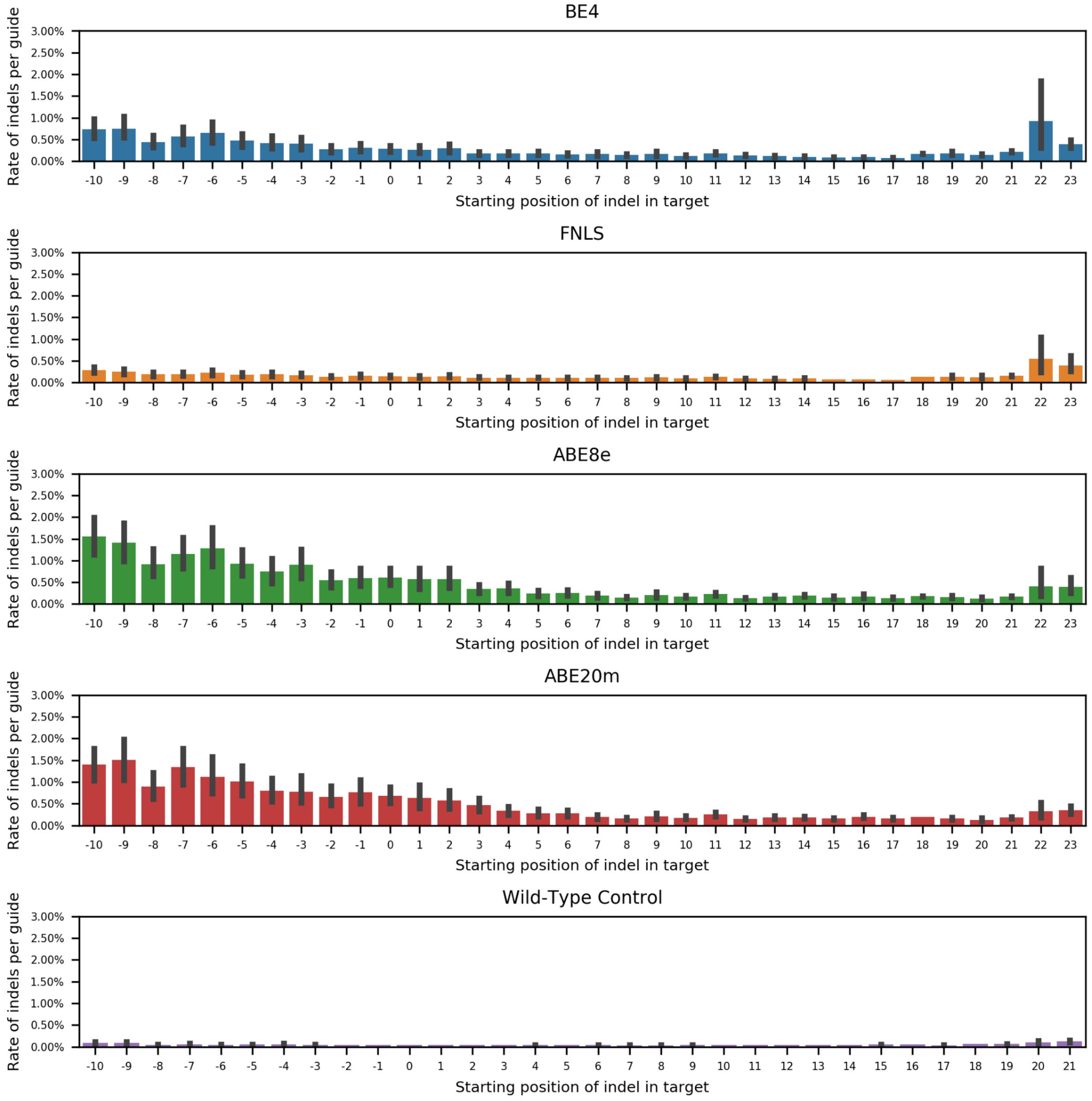


Insertions and deletion frequencies are low. Median frequency of indels per guide (y-axis) at each position in the target sequence (x-axis) for BE4 (top row), FNLS (second row), ABE8e (thrid row), ABE20m (fourth row) and cells without editors (bottom row). Error bars: 95% bootstrap confidence intervals from 1000 samples.

#### Figure S1I

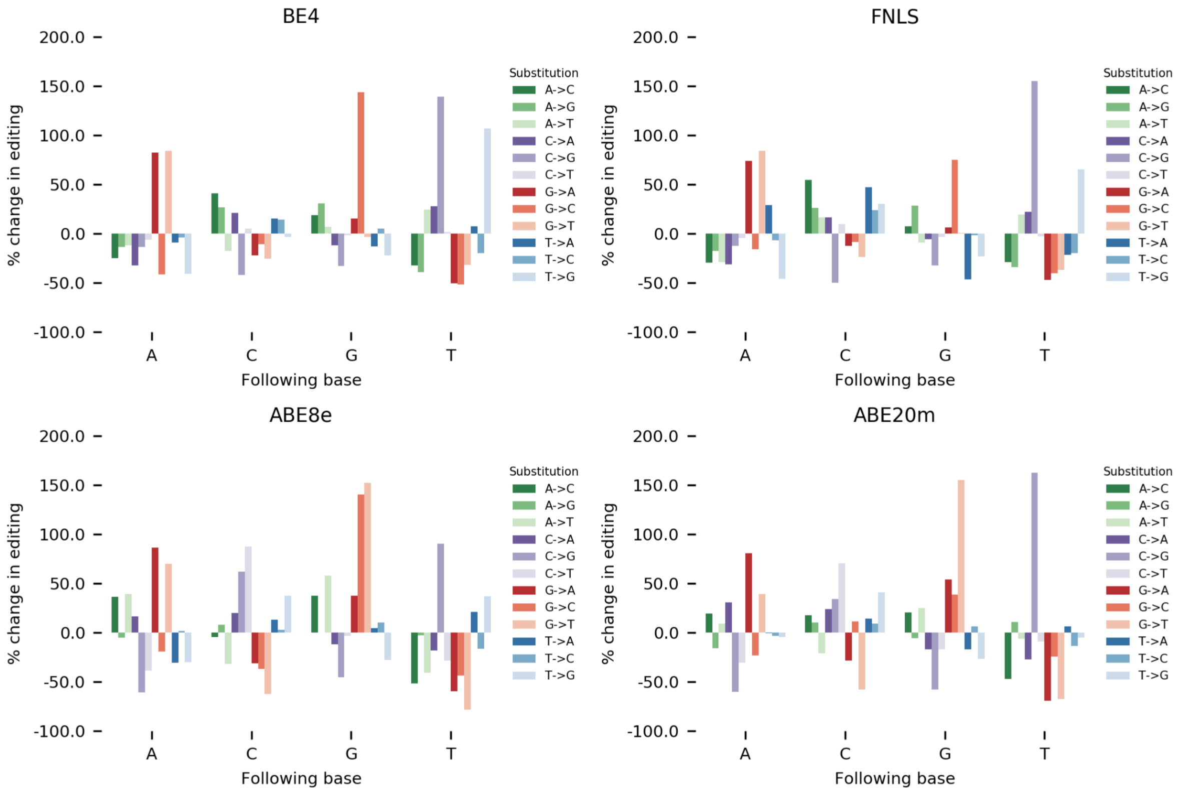


The influence of the base following the edited cytosine/adenine. Percentage change in editing rate (y-axis) when the target base is preceded by a certain base (x-axis) versus all other bases for BE4 (top left), FNLS (top right), ABE8e (bottom left) and ABE20m (bottom right). Colors: substitutions.

Figure S1J

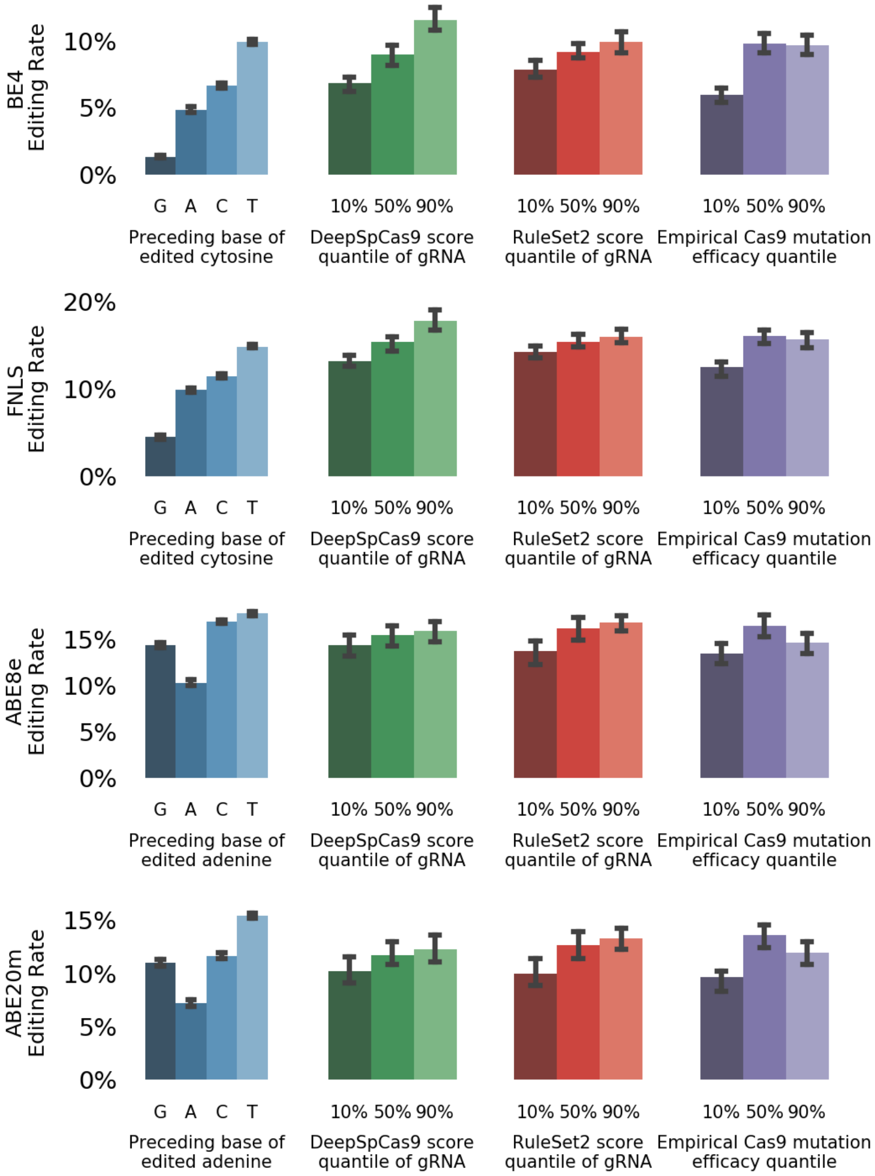


Cas9 gRNA features are informative of base editing rate. Median base editing rate (y-axis) is influenced by preceding base (blue bars), DeepSpCas9 score decile (green bars), RuleSet2 scores (red bars) and empirical Cas9 mutation efficacy (purple bars). Plots shown for BE4 (top row), FNLS (second row), ABE8e (third row) and ABE20m (bottom row). Error bars: 95% confidence intervals from 1000 bootstrap samples.

### Figure S2A

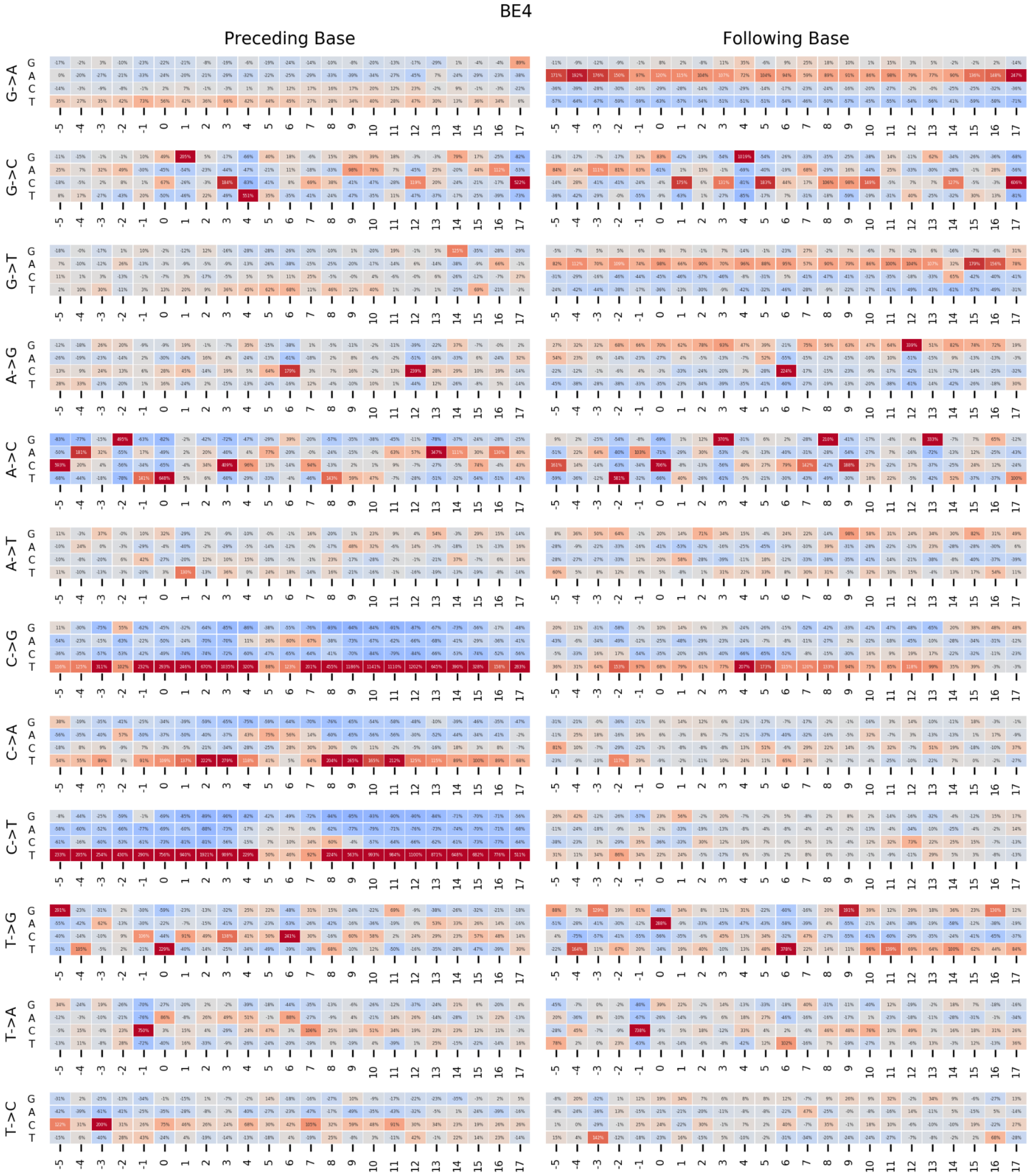


Position-dependent effect of the preceding and following base in BE4. Percentage change in editing rate (color) depending on the identity of the base (y-axis) preceding (left) or following (right) the target base in the sequence (x-axis) for each substitution type (rows).

### Figure S2B

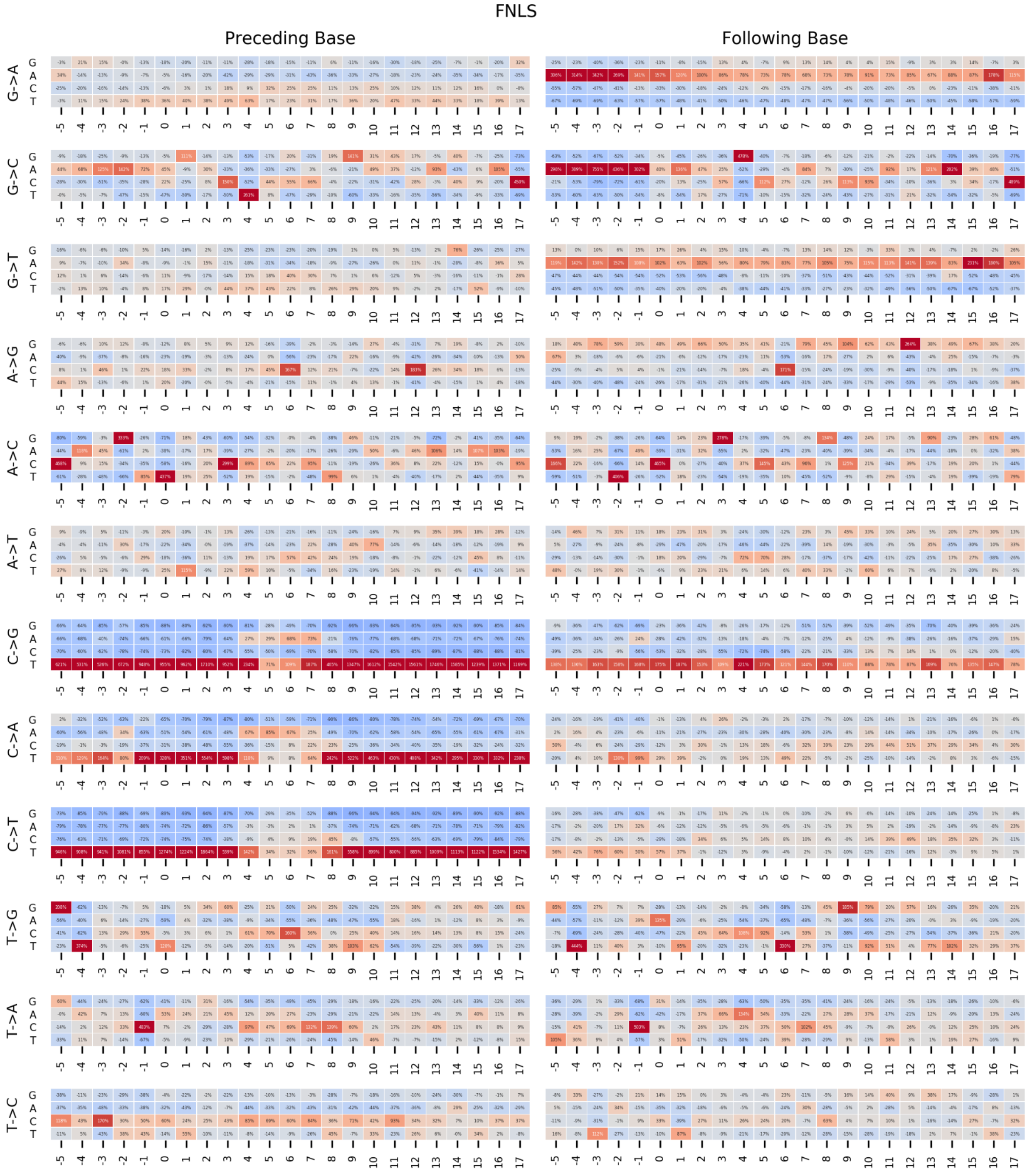


Position-dependent effect of the preceding and following base in FNLS. Percentage change in editing rate (color) depending on the identity of the base (y-axis) preceding (left) or following (right) the target base in the sequence (x-axis) for each substitution type (rows).

### Figure S2C

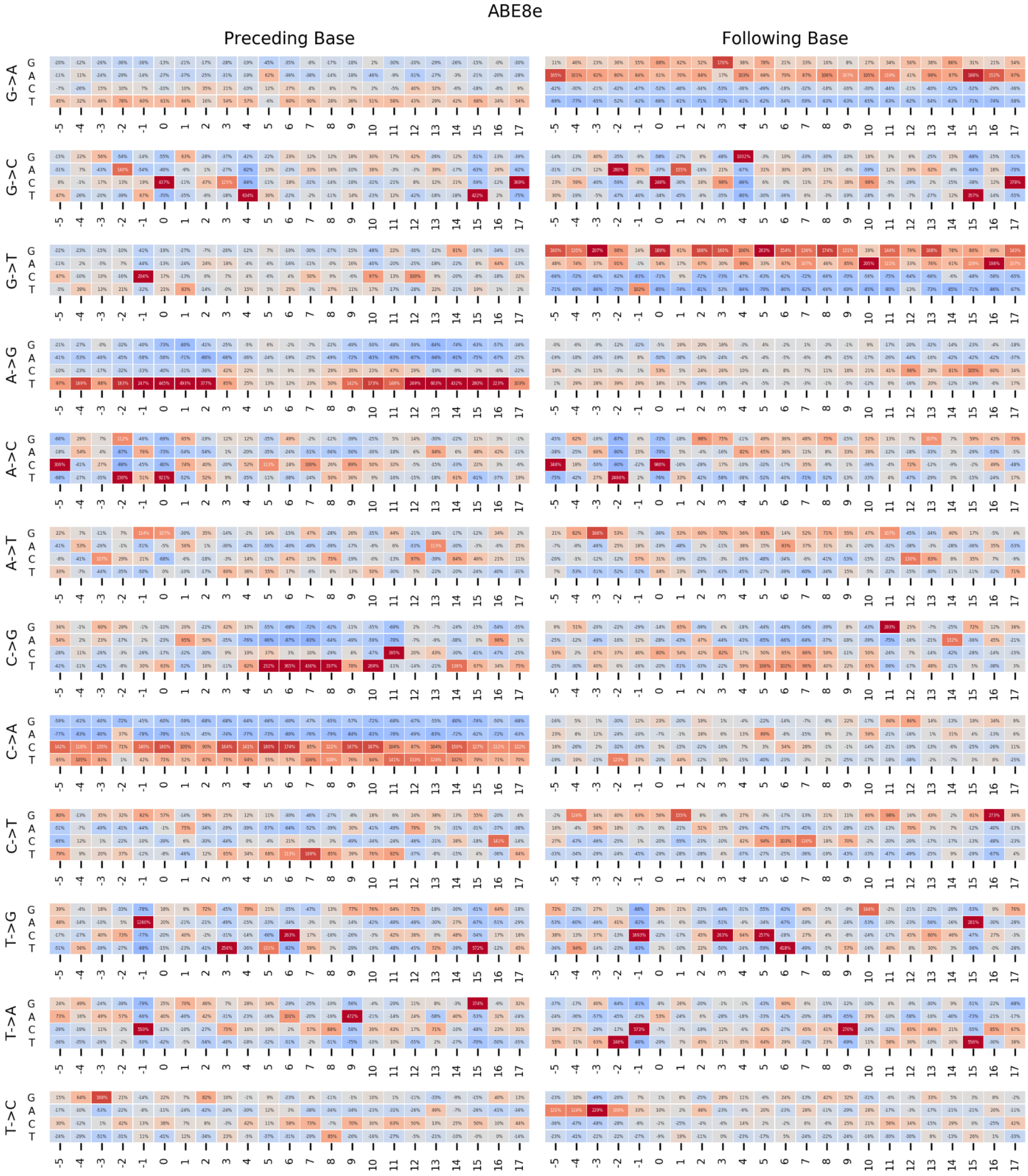


Position-dependent effect of the preceding and following base in ABE8e. Percentage change in editing rate (color) depending on the identity of the base (y-axis) preceding (left) or following (right) the target base in the sequence (x-axis) for each substitution type (rows).

Figure S2D

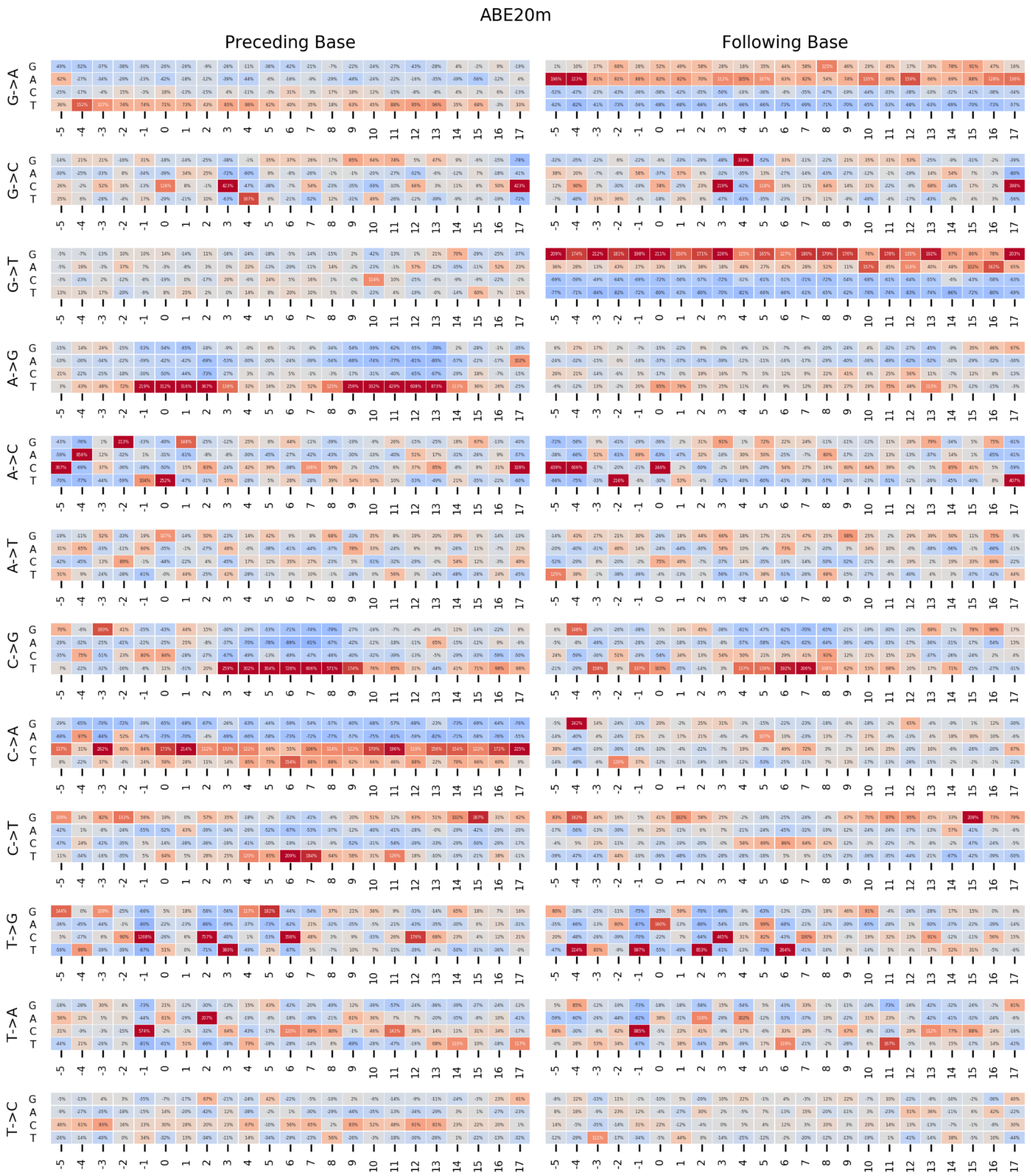


Position-dependent effect of the preceding and following base in ABE20m. Percentage change in editing rate (color) depending on the identity of the base (y-axis) preceding (left) or following (right) the target base in the sequence (x-axis) for each substitution type (rows).

### Figure S2E

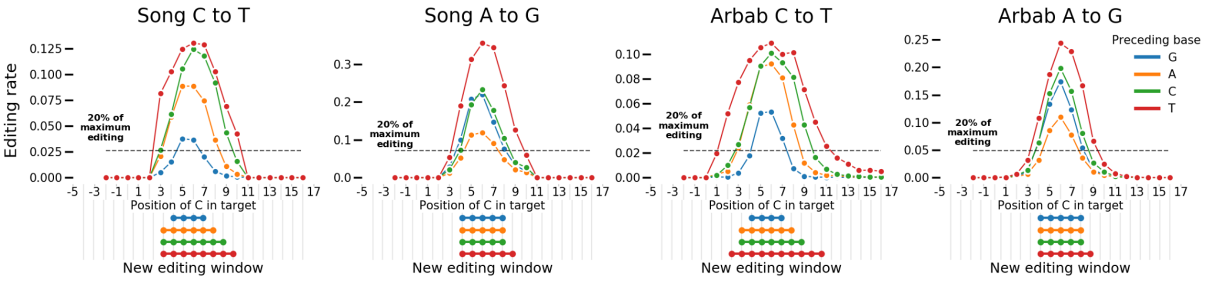


Window of editing changes depending on the base preceding the edited cytosine in other studies. Median editing rate (y-axis) of cytosines at positions -5 to 17 in the target (x-axis) for each preceding base type (colors) in datasets from Song et al (left) and Arbab et al (right) for each editor type (panels). Black dashed line: 20% of the maximum editing rate at any position for all preceding bases. Linked dots: positions at which editing is above 20% of the maximum.

### Figure S2F

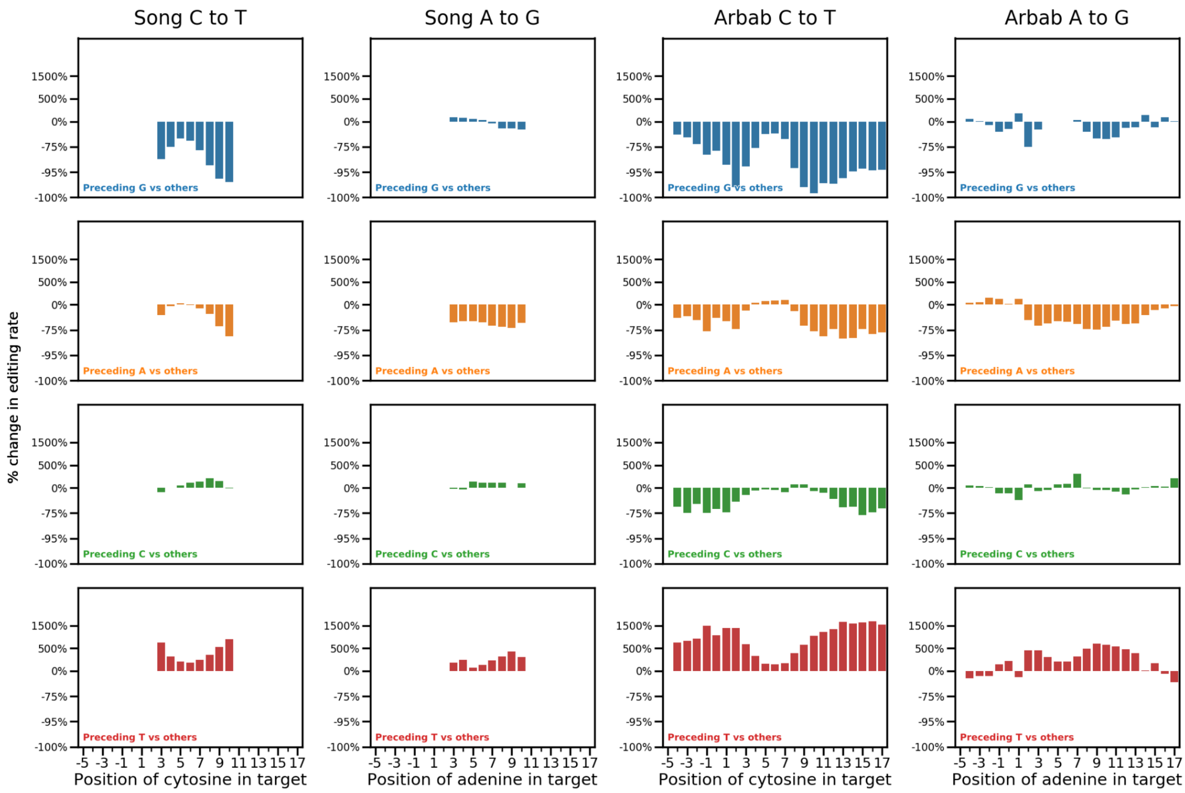


Position-dependent effect of the preceding base in Arbab and Song datasets for each type of editor (columns). Percentage change in editing rate (y-axis) from having a certain base preceding the cytosine (colors) compared to all other bases at different positions in the target sequence (x-axis).

#### Figure S2G

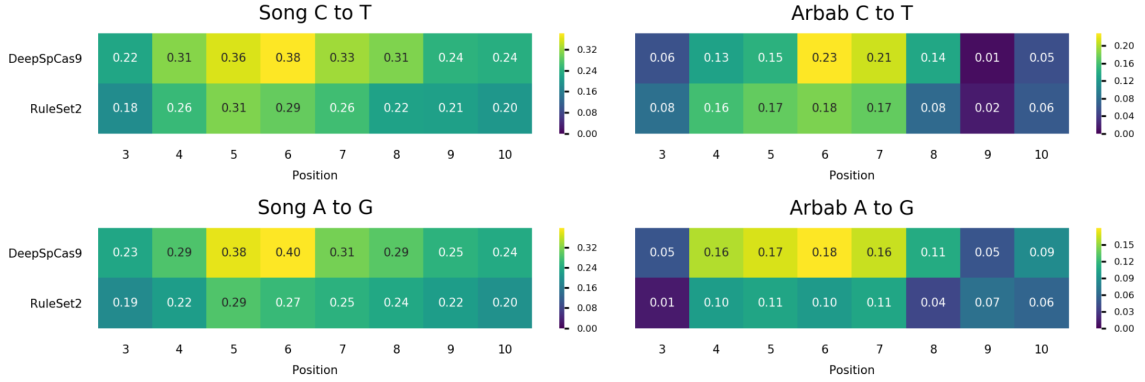


Correlation between gRNA quality and editing rate depend on the position in Song and Arbab datasets for each type of editor (panels). Pearson's R (color) between measures of gRNA quality (y-axis) and the position of the cytosine in the target sequence (x-axis).

Figure S3A

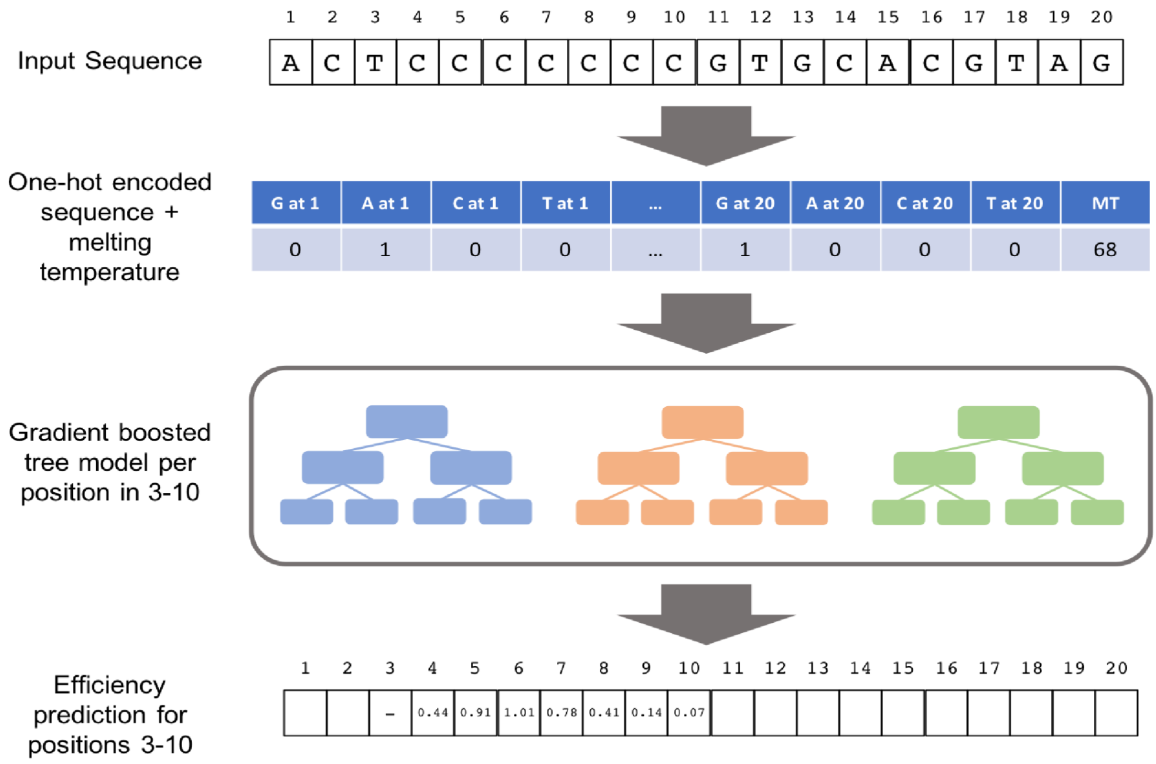


Outline of FORECasT-BE architecture. A 20 nucleotide input sequence is one-hot encoded into a feature vector and the melting temperature of the sequence is appended. A gradient boosted tree predictor uses these input features to predict a z-score representing editing efficiency for a given position in the target sequence. We train one gradient boosted tree predictor for each of positions 3-10.

### Figure S3B

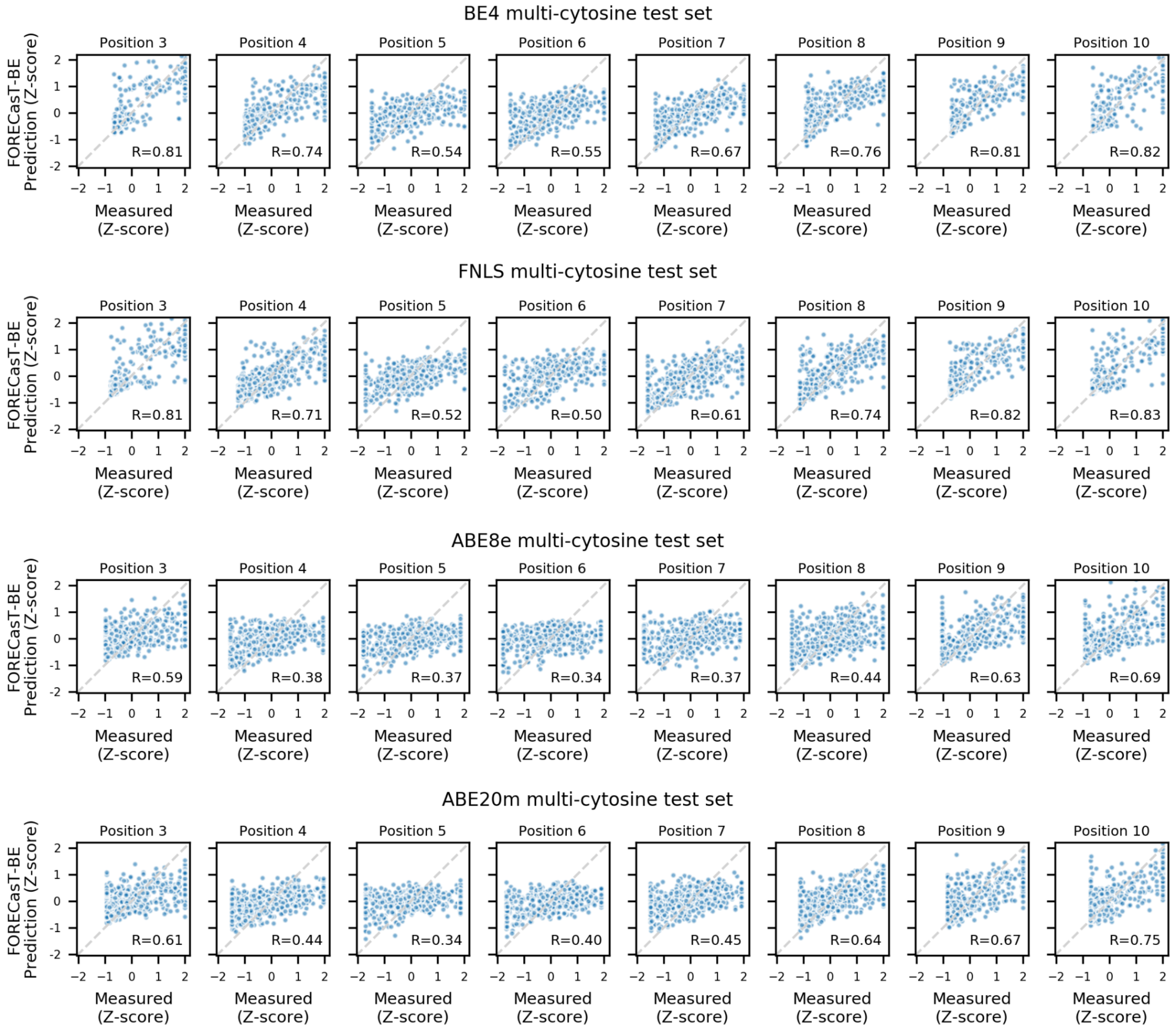


FORECasT-BE accurately predicts editing rate in multiple editors. Measured (x-axis) and predicted (y-axis) standardized editing rate (Methods) for guide RNAs (markers) with an adenine or cytosine at the different editing window positions (panels) for BE4 (top row), FNLS (second row), ABE8e (third row) and ABE20m (bottom row). Dashed line: y=x. Label: Pearson's R between measured and predicted scores.

### Figure S3C

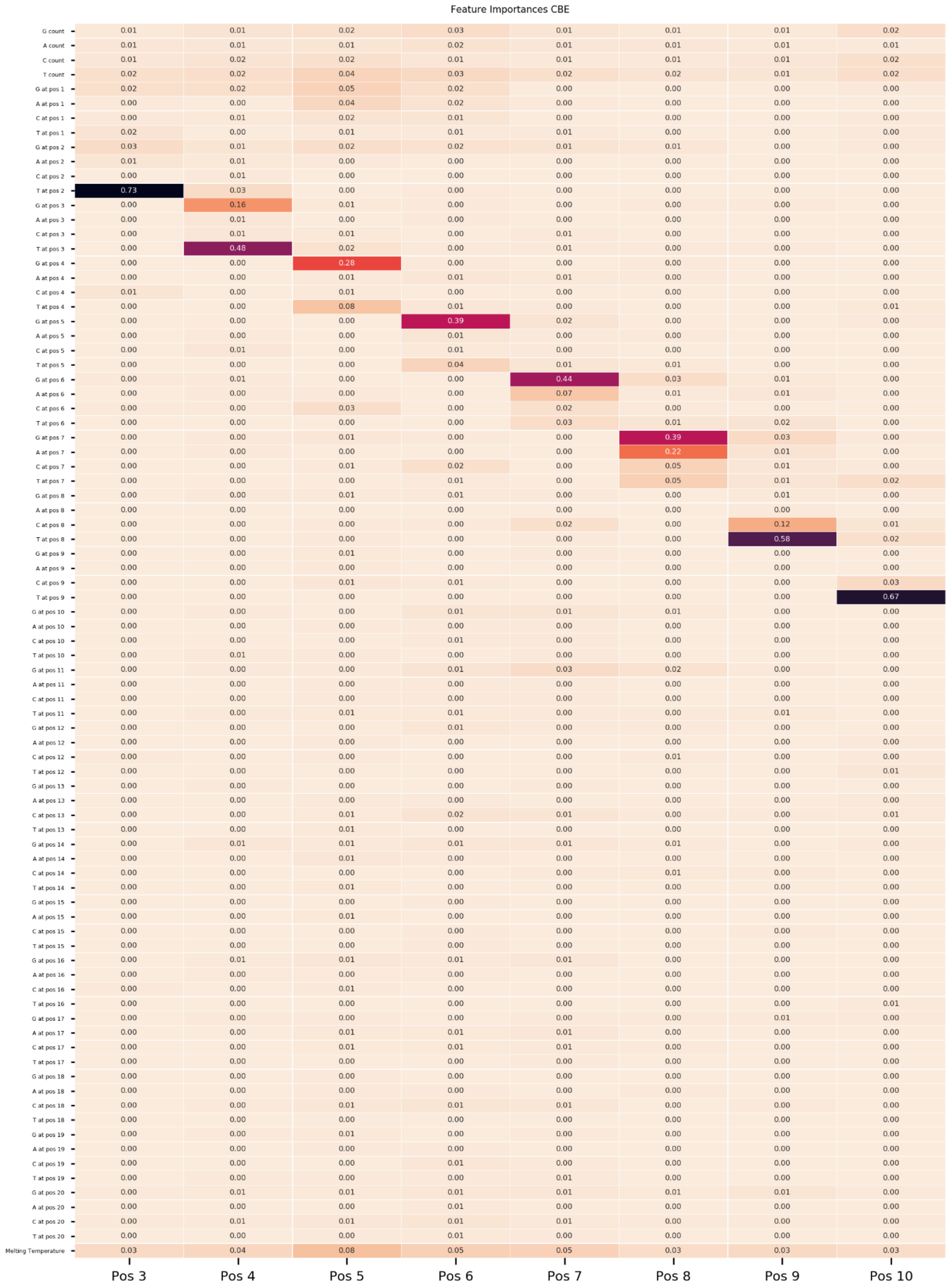


Feature importances for cytosine editors from FORECasT-BE. Gini coefficient (color) for every feature (y-axis) for each positional predictor (x-axis).

### Figure S3D

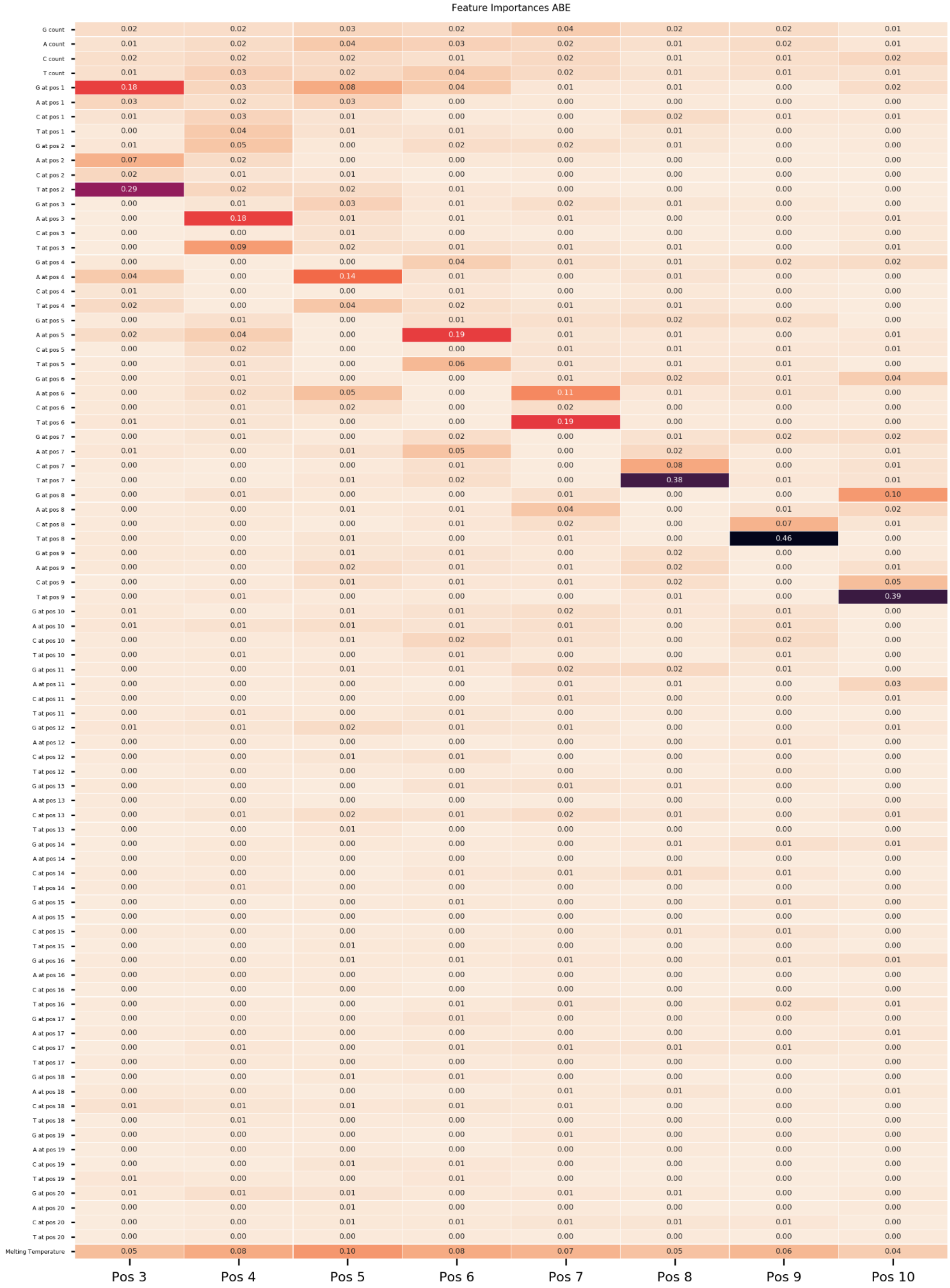


Feature importances for adenine editors from FORECasT-BE. Gini coefficient (color) for every feature (y-axis) for each positional predictor (x-axis).

#### Figure S3E

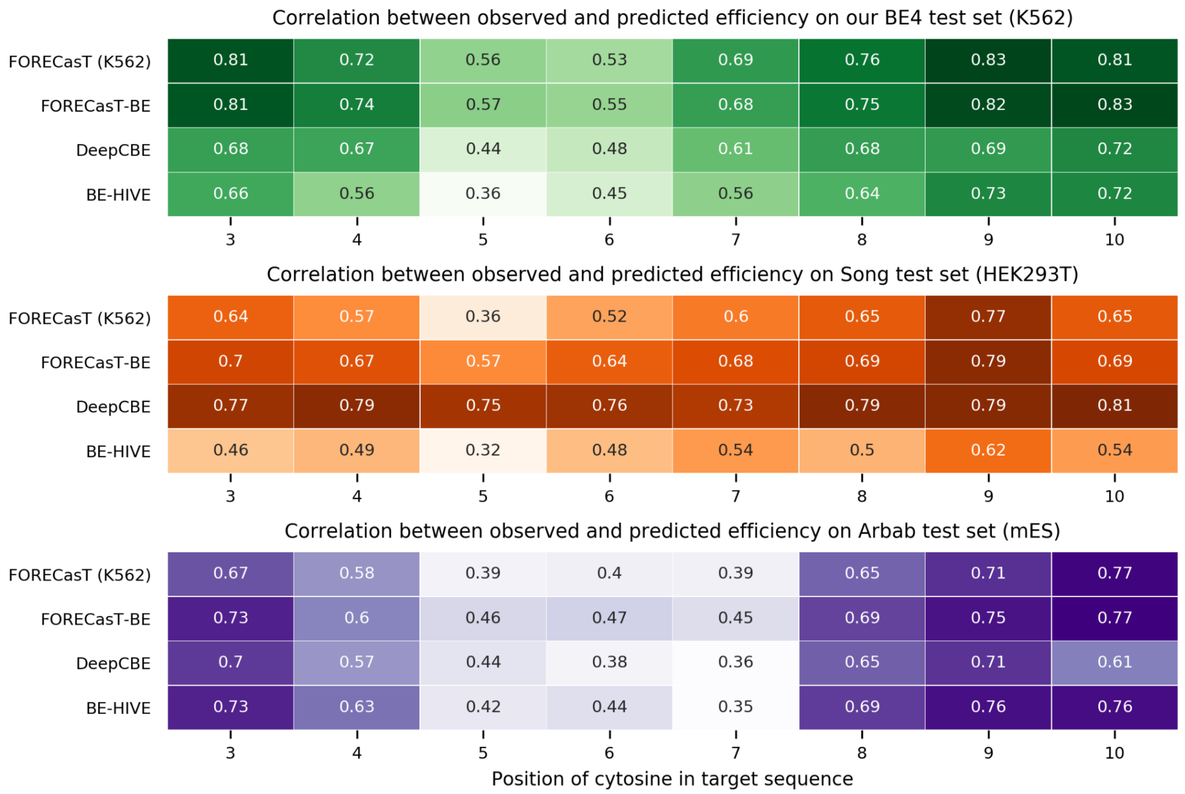


Cell type plays a part in prediction accuracy. Pearson's R between measurements and predictions for cytosine editor models trained on different cell types (y-axis) when using datasets with a cytosine at a given position in the guide sequence (x-axis). Models are evaluated on datasets from this study (K562, top heatmap), Song et al (HEK293T, middle) and Arbab et al (mES, bottom). Models compared on each dataset are: a FORECasT model trained on only K562 cells (top row), FORECasT-BE trained on multiple cell types (second row), DeepCBE trained on HEK293T cells (third row) and BE-HIVE trained on mES cells (bottom row).

### Figure S3F

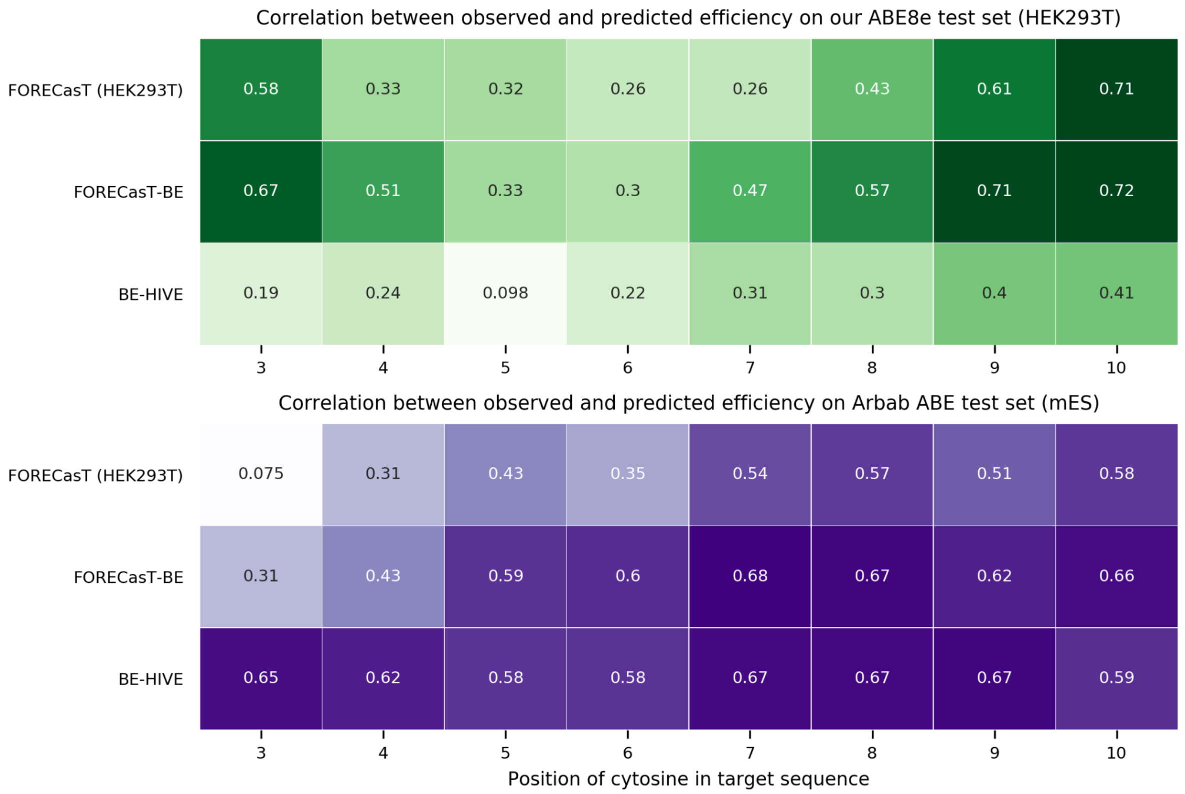


Cell type plays a part in prediction accuracy. Pearson's R between measurements and predictions for adenine editor models trained on different cell types (y-axis) when using datasets with an adenine at a given position in the guide sequence (x-axis). Models are evaluated on datasets from this study (HEK293T, top heatmap) and Arbab et al (mES, bottom). Models compared on each dataset are: a FORECasT model trained on only our HEK293T cells (top row), FORECasT-BE trained on multiple cell types (second row), and BE-HIVE trained on mES cells (bottom row).

### Figure S3G

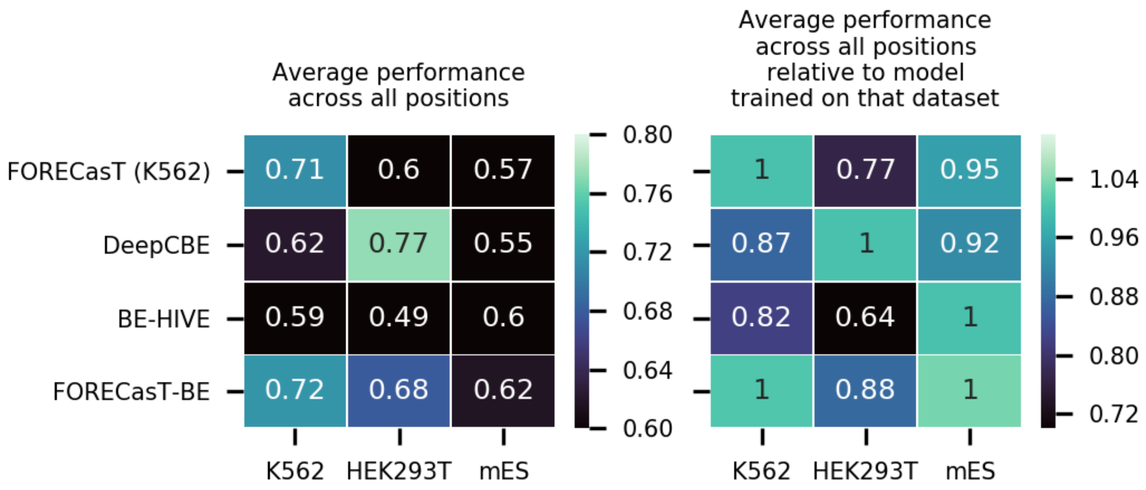


FORECasT-BE generalizes well to many cytosine datasets. Average performance of models (y-axis) at all positions when evaluated on a test set from a certain cell type (x-axis), both raw (left) and normalized (right). Normalization is relative to the performance of the model trained on that dataset (eg: FORECasT-K562 was trained on K562 cells so has a value of 1, while all other values in the column are a fraction of the performance of FORECasT-K562). Models compared are: FORECasT model trained only on K562 cells (top row), DeepCBE trained on HEK293T cells (second row), BE-HIVE trained on mES cells (third row) and FORECasT-BE model trained on multiple cell types (bottom row).

#### Figure S3H

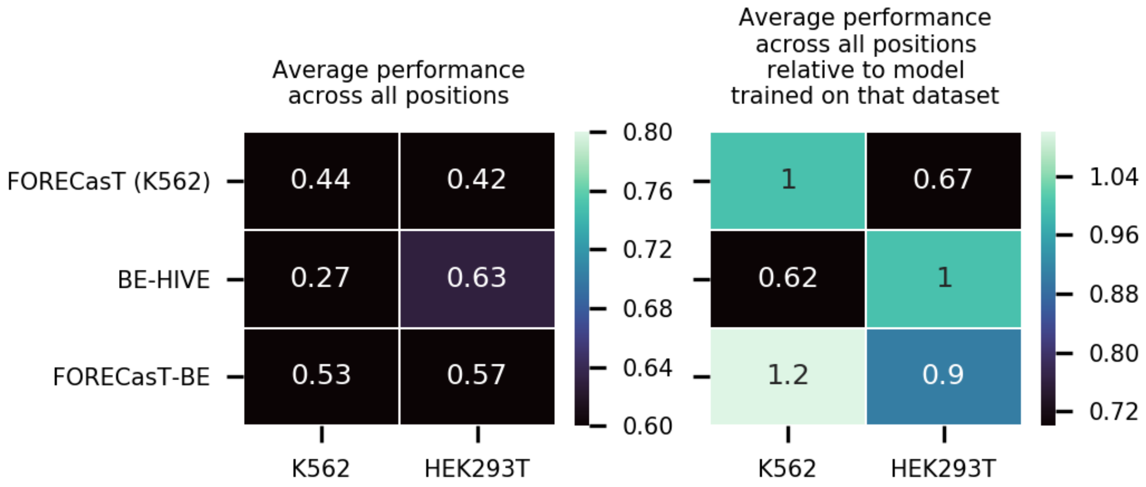


FORECasT-BE generalizes well to adenine datasets. Average performance of models (y-axis) at all positions when evaluated on a test set from a certain cell type (x-axis), both raw (left) and normalized (right). Normalization is relative to the performance of the model trained on that dataset (eg: FORECasT-K562 was trained on K562 cells so has a value of 1, while all other values in the column are a fraction of the performance of FORECasT-K562). Models compared are: FORECasT model trained only on K562 cells (top row), BE-HIVE trained on mES cells (second row) and FORECasT-BE model trained on multiple cell types (bottom row).

#### Figure S3I

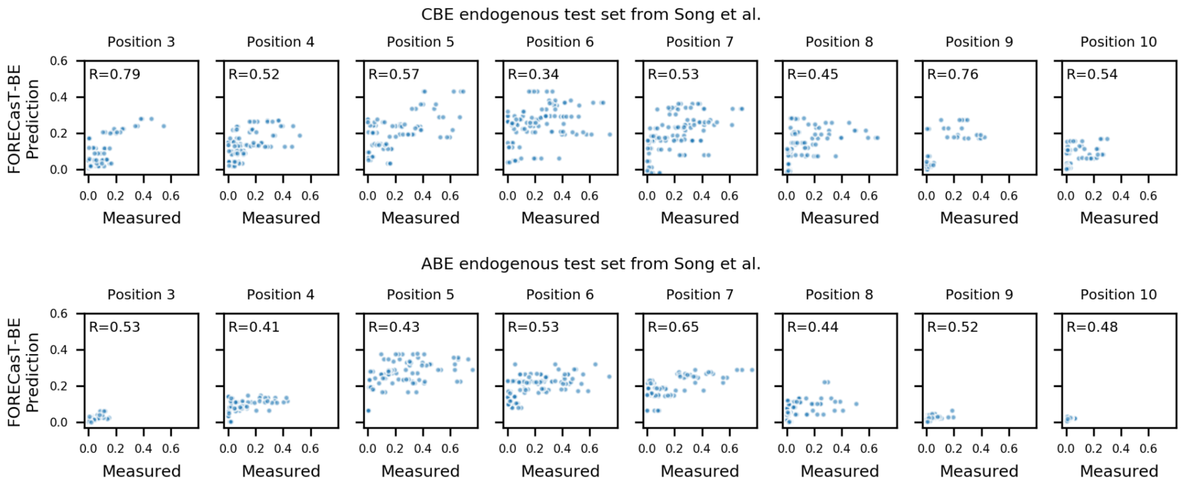


FORECasT-BE accurately predicts editing rate in an endogenous context [20]. Measured (x-axis) and predicted (y-axis) standardized editing rate (Methods) for guide RNAs (markers) with a cytosine (top) or adenine (bottom) at the different editing window positions (panels). Dashed line: y=x. Label: Pearson's R between the measured and predicted scores.

#### Figure S3J

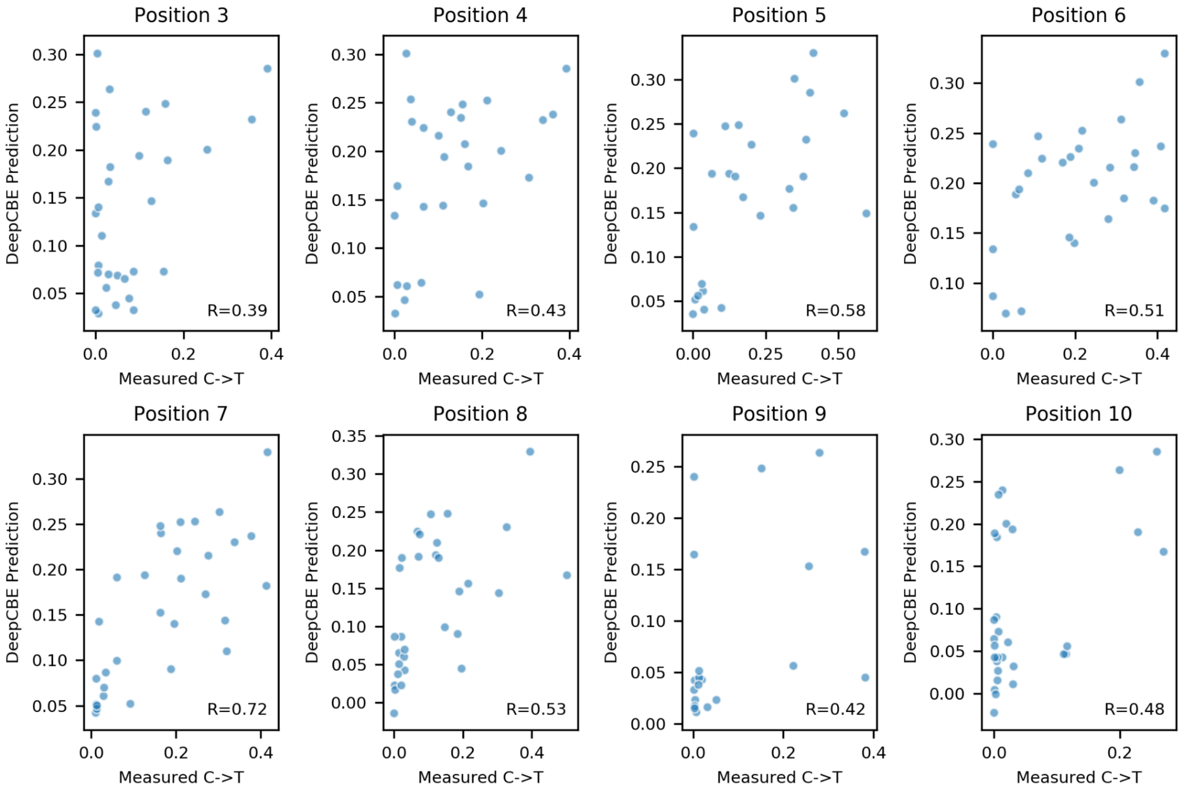


Predictive performance of DeepCBE on endogenous dataset from Song et al.. Comparison of predicted editing efficiency (y-axis) against measured editing efficiency (x-axis) for guides with cytosines at positions 3-10 (panels). Dashed line: y=x. Label: Pearson's R between measured and predicted scores

Figure S3K

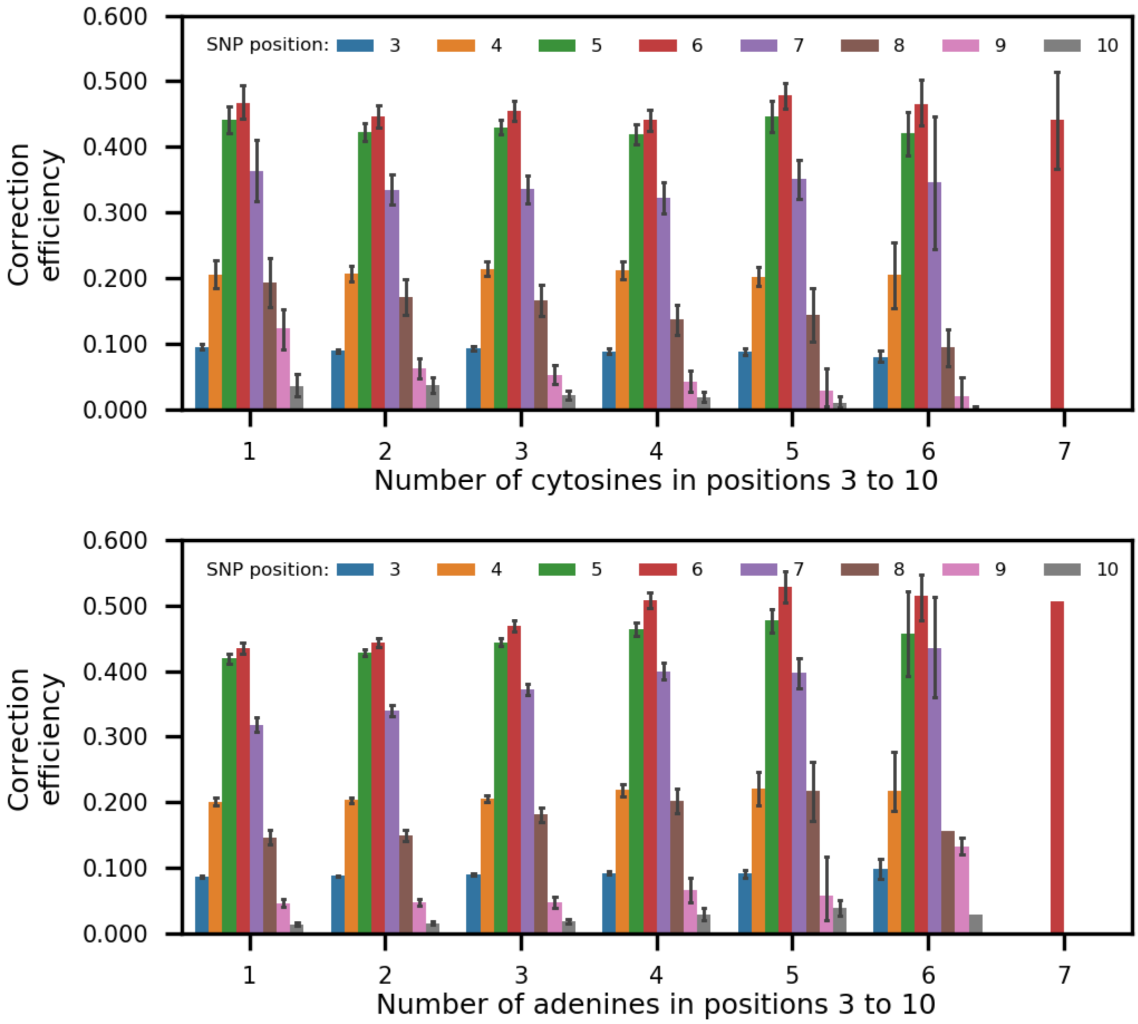


Correction efficiency varied across guides designed to correct pathogenic SNPs. Predicted efficiency to correct a pathogenic SNP (y-axis), for guides with differing numbers of cytosines (top) or adenines (bottom) in positions 3-10 (x-axis), and having SNPs at different positions in their sequence (colors). Error bars: 95% confidence intervals from 1000 bootstrap samples.

### Figure S4A

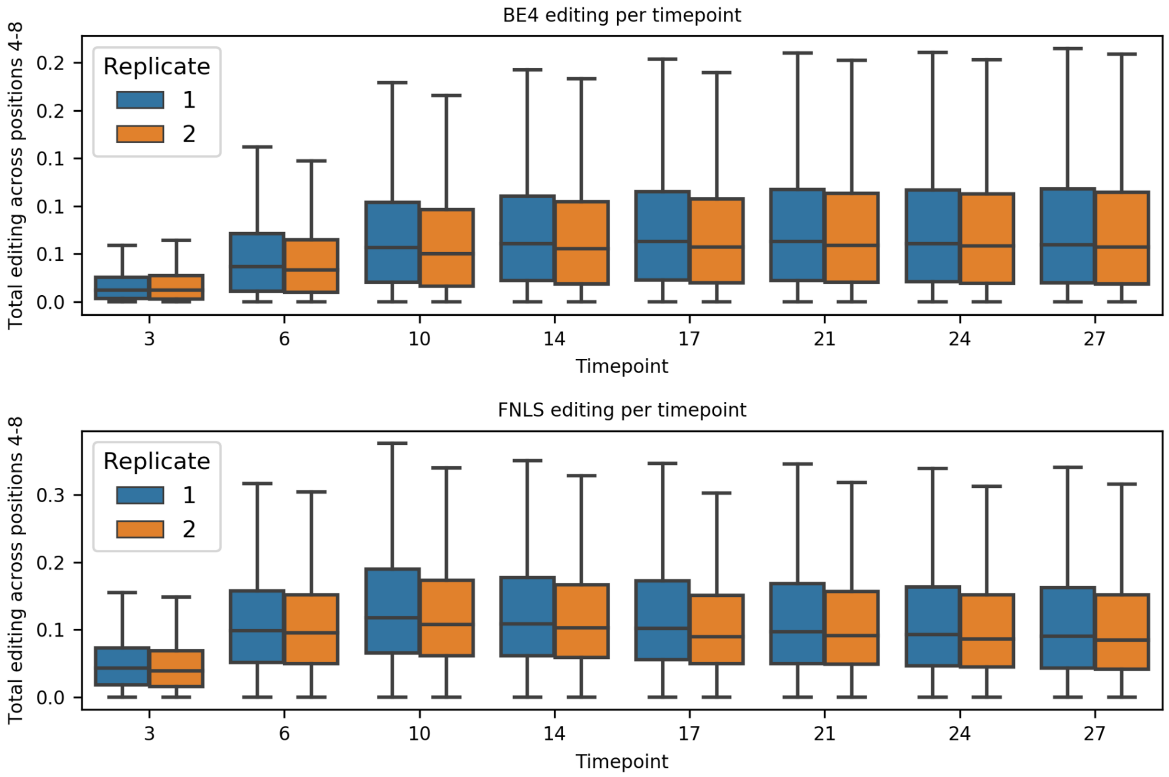


Editing plateaus at day 10, but slowly decreases afterward. Total editing measured across positions 4-8 (y-axis) for samples taken at each timepoint after infection in the screen (x-axis), for each replicate (colors). Boxes: median and quartiles; whiskers: 1.5 interquartile ranges from the top and bottom quartiles (bounded by 0 and 1)

### Figure S4B

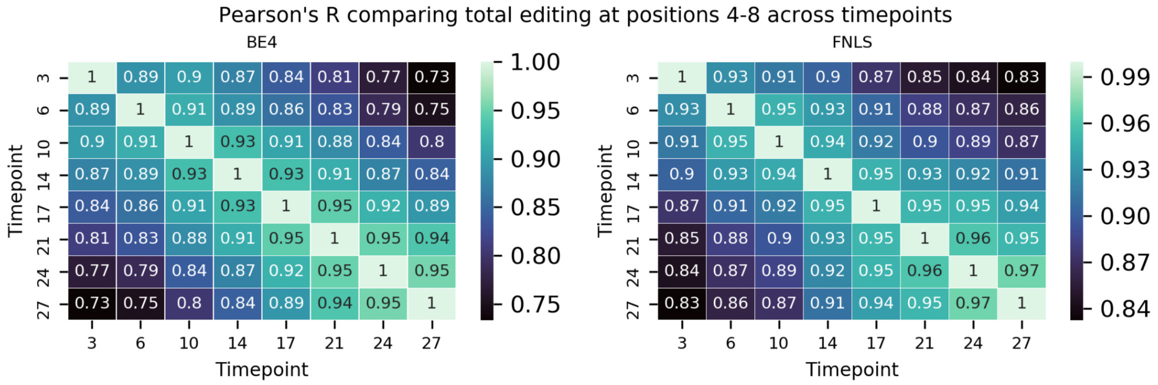


Closer timepoints edit similarly. Pearson’s R between total editing at positions 4-8 at different timepoints (x- and y-axes) .

#### Figure S4C

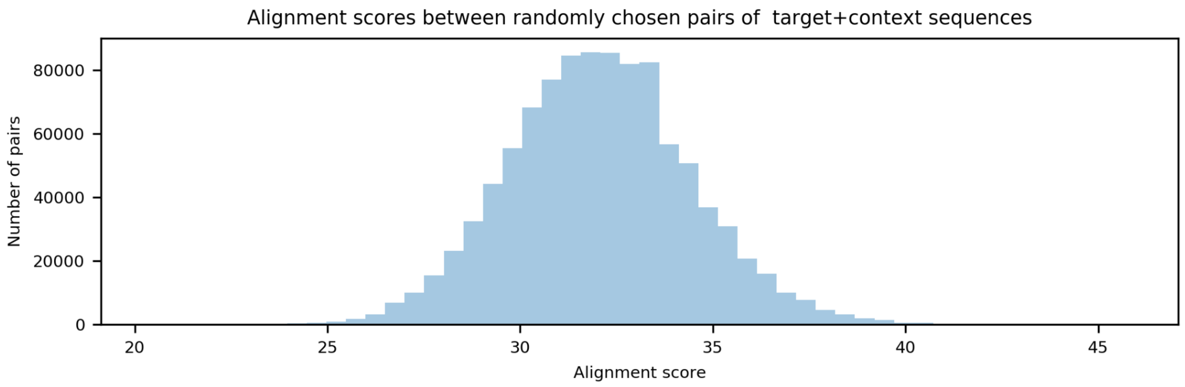


Alignment scores between randomly chosen 79nt target+context sequences in our constructs. Number of randomly chosen pairs of 79nt target+context sequences (y-axis) with a specific alignment score (x-axis).

#### Table S1

| Primer | Sequence |
|---|---|
| P1 | ACACTCTTTCCCTACACGACGCTCTTCCGATCTGACGTCCAGAGCACAGATGG |
| P2 | TCGGCATTCCTGCTGAACCGCTCTTCCGATCTACCCGGTAGAATTGGATCCAAAC |
| P3 | AATGATACGGCGACCACCGAGATCTACACTCTTTCCCTACACGACGCTCTTCCGATC*T |
| P4* | CAAGCAGAAGACGGCATACGAGAT$N_{10}$GAGATCGGTCTCGGCATTCCTGCTGAACCGCTCTTCCGATC T |
| P5 | ACACTCTTTCCCTACACGACGCTCTTCCGATCTCTTGGCTTTATATATCTTGTGGAAAGGACGAAACA |
| P6 | TCGGCATTCCTGCTGAACCGCTCTTCCGATCTCTAAAGCGCATGCTCCAGACTGCC |

*index for multiplexed sequencing

#### Table S2

| SANGER SAMPLE ID | SAMPLE LABEL | ERS | SAMPLE DESCRIPTION |
|---|---|---|---|

| | | | |
|---|---|---|---|
| T227_CRISPR8015850 | 22_1_3D14 | ERS3536251 | ABERA Replicate 1 Day 14 |
| T227_CRISPR8015851 | 22_1_3D17A | ERS3536252 | ABERA Replicate 1 Day 17A |
| T227_CRISPR8015852 | 22_1_3D17B | ERS3536253 | ABERA Replicate 1 Day 17B |
| T227_CRISPR8015853 | 22_1_3D17C | ERS3536254 | ABERA Replicate 1 Day 17C |
| T227_CRISPR8015854 | 22_1_3D17D | ERS3536255 | ABERA Replicate 1 Day 17D |
| T227_CRISPR8015855 | 22_1_3D21 | ERS3536256 | ABERA Replicate 1 Day 21 |
| T227_CRISPR8015856 | 22_1_3D24 | ERS3536257 | ABERA Replicate 1 Day 24 |
| T227_CRISPR8015857 | 22_1_3D27 | ERS3536258 | ABERA Replicate 1 Day 27 |
| T227_CRISPR8015858 | 22_1_4D14 | ERS3536259 | ABERA Replicate 2 Day 14 |
| T227_CRISPR8015859 | 22_1_4D21 | ERS3536260 | ABERA Replicate 2 Day 21 |
| T227_CRISPR8015860 | 22_1_4D24 | ERS3536261 | ABERA Replicate 2 Day 24 |
| T227_CRISPR8015861 | 22_1_4D27 | ERS3536262 | ABERA Replicate 2 Day 27 |
| T227_CRISPR8015862 | 22_1_5D3 | ERS3536263 | BE4 Replicate 1 Day 3 |
| T227_CRISPR8015863 | 22_1_5D6 | ERS3536264 | BE4 Replicate 1 Day 6 |
| T227_CRISPR8015864 | 22_1_5D10 | ERS3536265 | BE4 Replicate 1 Day 10 |
| T227_CRISPR8015865 | 22_1_5D14 | ERS3536266 | BE4 Replicate 1 Day 14 |
| T227_CRISPR8015866 | 22_1_5D17 | ERS3536267 | BE4 Replicate 1 Day 17 |
| T227_CRISPR8015867 | 22_1_5D21 | ERS3536268 | BE4 Replicate 1 Day 21 |
| T227_CRISPR8015868 | 22_1_5D24 | ERS3536269 | BE4 Replicate 1 Day 24 |
| T227_CRISPR8015869 | 22_1_5D27 | ERS3536270 | BE4 Replicate 1 Day 27 |
| T227_CRISPR8015870 | 22_1_6D3 | ERS3536271 | BE4 Replicate 2 Day 3 |
| T227_CRISPR8015871 | 22_1_6D6 | ERS3536272 | BE4 Replicate 2 Day 6 |
| T227_CRISPR8015872 | 22_1_6D10 | ERS3536273 | BE4 Replicate 2 Day 10 |
| T227_CRISPR8015873 | 22_1_6D14 | ERS3536274 | BE4 Replicate 2 Day 14 |
| T227_CRISPR8015874 | 22_1_6D17 | ERS3536275 | BE4 Replicate 2 Day 17 |

| | | | |
|---|---|---|---|
| T227_CRISPR8015875 | 22_1_6D21 | ERS3536276 | BE4 Replicate 2 Day 21 |
| T227_CRISPR8015876 | 22_1_6D24 | ERS3536277 | BE4 Replicate 2 Day 24 |
| T227_CRISPR8015877 | 22_1_6D27 | ERS3536278 | BE4 Replicate 2 Day 27 |
| T227_CRISPR8015878 | 22_1_7D3 | ERS3536279 | FNLS Replicate 1 Day 3 |
| T227_CRISPR8015879 | 22_1_7D6 | ERS3536280 | FNLS Replicate 1 Day 6 |
| T227_CRISPR8015880 | 22_1_7D10 | ERS3536281 | FNLS Replicate 1 Day 10 |
| T227_CRISPR8015881 | 22_1_7D14 | ERS3536282 | FNLS Replicate 1 Day 14 |
| T227_CRISPR8015882 | 22_1_7D17 | ERS3536283 | FNLS Replicate 1 Day 17 |
| T227_CRISPR8015883 | 22_1_7D21 | ERS3536284 | FNLS Replicate 1 Day 21 |
| T227_CRISPR8015884 | 22_1_7D24 | ERS3536285 | FNLS Replicate 1 Day 24 |
| T227_CRISPR8015885 | 22_1_7D27 | ERS3536286 | FNLS Replicate 1 Day 27 |
| T227_CRISPR8015886 | 22_1_8D3 | ERS3536287 | FNLS Replicate 2 Day 3 |
| T227_CRISPR8015887 | 22_1_8D6 | ERS3536288 | FNLS Replicate 2 Day 6 |
| T227_CRISPR8015888 | 22_1_8D10 | ERS3536289 | FNLS Replicate 2 Day 10 |
| T227_CRISPR8015889 | 22_1_8D14 | ERS3536290 | FNLS Replicate 2 Day 14 |
| T227_CRISPR8015890 | 22_1_8D17 | ERS3536291 | FNLS Replicate 2 Day 17 |
| T227_CRISPR8015891 | 22_1_8D21 | ERS3536292 | FNLS Replicate 2 Day 21 |
| T227_CRISPR8015892 | 22_1_8D24 | ERS3536293 | FNLS Replicate 2 Day 24 |
| T227_CRISPR8015893 | 22_1_8D27 | ERS3536294 | FNLS Replicate 2 Day 27 |
